# AI-assisted Image-Based Phenotyping Reveals Genetic Architecture of Pod Traits in Mungbean (*Vigna radiata* L.)

**DOI:** 10.1101/2025.11.06.687031

**Authors:** Venkata Naresh Boddepalli, Talukder Zaki Jubery, Somak Dutta, Baskar Ganapathysubramanyan, Arti Singh

## Abstract

Mungbean (*Vigna radiata* (L.) R. Wilczek) is a vital source of digestible proteins and is well-suited for the plant-based protein industry. In this study, we analyzed pod morphological traits in the Iowa Mungbean Diversity (IMD) panel with 372 genotypes (2022-23) with AI-assisted image phenotyping using 2,418 pod images. Pod morphological traits were extracted using deep learning image analysis, achieving excellent agreement with manual measurements (r>0.96 for pod length and seed per pod). Four complementary GWAS models identified 45 significant SNPs associated with pod curvature, length, width, and seed per pod traits. Notably, a significant SNP (5_35265704) on chromosome 1 was linked to pod dimensional traits, length, width, and curvature. A candidate gene, *Vradi01g00001116*, was located within the linkage disequilibrium (LD) region of this SNP, is part of the GH3 gene family, and has an *Arabidopsis* ortholog (*AT4G27260*) known for influencing organ elongation, pod, and seed development. Another SNP, 5_210437 on chromosome 2, has been found to be significantly associated with both pod length and seed per pod. A candidate gene, *Vradi02g00003971*, located in the LD region of this SNP, belongs to the potassium transporter family and shares homology with the HAK5 gene family (*AT4G13420*) in *Arabidopsis*, which influences pod and seed growth. Image-based measurements achieved genomic prediction accuracies ranging from 0.61 to 0.85 across various traits, exhibiting an improvement of 12-22% over manual methods. These results demonstrate the potential of AI-assisted phenomics integrated with genomic tools to accelerate selection for improved pod architecture in mungbean breeding programs across the Midwestern United States and globally.

## 1. INTRODUCTION

Mungbean (*Vigna radiata* (L.) R. Wilczek) is a warm-season food grain legume crop globally known for its high nutritional value. With over 7 million hectares of global cultivated area, predominantly grown in Asian countries, followed by Australia, several African nations, South America, and parts of the USA. (Kohno et al. 2018; Nair and Schreinemachers 2020; Somta et al. 2022; Fumia et al. 2023). Mungbean seeds contain 20-30% digestible proteins, dietary fiber, essential amino acids, iron, and folates (Nair and Schreinemachers 2020; Sandhu and Singh 2021; Somta et al. 2022). Mungbean has historically been a staple protein source in South and Southeast Asia (Somta et al. 2022). In recent years, mungbean has emerged as a promising option for crop diversification in North America, including the United States and Canada, where there is a growing demand for sustainable plant-based protein sources (Sandhu and Singh 2021). Its ability to thrive in hot, drought-prone conditions, with a short lifecycle (60–75 days after planting), and most importantly, its ability to share the same infrastructure needed for soybean cultivation with minimal adjustments, make it a valuable addition to existing cropping systems. Despite its global importance, mungbean remains underutilized and underrepresented in U.S. agricultural systems and breeding programs (Sandhu and Singh 2021). However, recent initiatives, such as the establishment of the Iowa Mungbean Diversity (IMD) panel, aim to address this gap by evaluating germplasm under the U.S. field conditions for agronomic suitability (Sandhu and Singh 2021). These efforts highlight the potential of mungbeans to not only supplement domestic protein needs but also enhance ecological and economic resilience through diversification.

In legumes, pods are essential reproductive structures that can directly affect yield through their morphological traits, including pod length, width, curvature, and the number of seeds per pod. These traits are potential determinants of harvest index and yield, also affecting threshability, seed uniformity, and market value (Funatsuki et al. 2014; Di Vittori et al. 2021; Li et al. 2024; Njau et al. 2024). During mungbean domestication, pod architecture and dehiscence were important traits, with favorable selection towards larger, non-shattering pods that contain more seeds per pod (Li et al. 2024). Previous studies documented a positive correlation between pod length and the number of seeds per pod with seed yield and 100-seed weight in mungbean (Liu et al. 2022; Chang et al. 2023; Manjunatha et al. 2023; Li et al. 2024), common bean (García-Fernández et al. 2021), cowpea (Xu et al. 2017), soybean (Chen et al. 2023), and broad bean (Li and Yang 2014). These studies highlight the need to understand the genetic architecture underlying these yield-contributing traits, where genome-wide association studies (GWAS) and comparative mapping enable the identification of major loci and candidate genes; however, there have been limited studies on mungbean pod morphological traits using large diversity panels and multi-environment phenotypic data (Liu et al., 2022; Somta et al., 2022; Manjunatha et al., 2023; Li et al., 2024). The complex quantitative nature of pod morphological traits, combined with their high heritability and breeding importance, makes them ideal candidates for genomic dissection using modern molecular approaches. However, understanding the genetic control of these traits is essential for marker-assisted selections that can accelerate breeding progress and enable precise manipulation of pod architecture in mungbean improvement programs.

Recent advancements in genomics have led to increased access to modern genetic tools, such as GWAS, comparative mapping, and genomic prediction (GP), for accelerating trait discovery and new cultivar development (Han et al. 2022; Somta et al. 2022; Fumia et al. 2023; Ahmed et al. 2024; Chiteri et al. 2024). GWAS has effectively identified quantitative trait loci (QTL) related to important agronomic traits in mungbean, such as 100-seed weight, flowering time, leaf and root morphology, pod dehiscence, biotic and abiotic resistance, using the high-density single nucleotide polymorphism (SNP) data from various mungbean panels (Chiteri et al., 2021; Han et al. 2022; Chiteri et al. 2023; Manjunatha et al. 2023; Ahmed et al. 2024; Chiteri et al. 2024; Kohli et al. 2024; Li et al. 2024; Iqbal et al. 2025). Comparative mapping with closely related legumes, such as cowpea (*Vigna unguiculata* (L.) Walp.), common bean (*Phaseolus vulgaris* L.), and soybean (*Glycine max* (L.) Merr.), has revealed conserved synteny and orthologous regions that regulate domestication traits. This enables researchers to understand gene functions across different species (Manjunatha et al. 2023; Chiteri et al. 2024). Moreover, genomic prediction models that include trait variations have shown great potential for increasing selection accuracy and reducing breeding cycles (Miller et al. 2023; Tang et al. 2024; Escamilla et al. 2025; Mbebi et al. 2025). To the best of our knowledge, no genomic prediction studies have been specifically published on mungbean, although there have been a few attempts recently with other legumes (Keller et al. 2020; Shao et al. 2022; Huynh et al. 2024; Van der Laan et al. 2025). By integrating phenotypic data from multiple environments with genome-wide marker effects, these models offer a scalable approach for enhancing genetic improvement in mungbean. However, precise phenotyping of pod morphological traits for genetic studies is a significant challenge in legumes. Traditional manual measurements are labor-intensive, time-consuming, and prone to inter-and intra-rater variability. This variability can diminish the statistical power of subsequent GWAS and GP models(Li et al. 2020; Chang et al. 2021; Gill et al. 2022; Yang et al. 2024; Liu et al. 2025).

To address these challenges in mungbean breeding, this study integrated high-throughput image analysis with genomic approaches to examine the genetic structure of pod morphological traits. Our study objectives were to: (1) develop and validate machine learning model for the automated extraction of pod length, width, curvature, and seed per pod traits from scanned images; (2) characterize the phenotypic diversity and trait relationships within the Iowa Mungbean Diversity panel; (3) identify the marker trait association of these traits through genome-wide association studies; and (4) evaluate the effectiveness of image-based phenotyping compared to manual phenotyping for genomic prediction. This integrated approach provides essential tools and genetic insights aimed at accelerating mungbean improvement through marker-assisted breeding and genomic selection.

## 2. MATERIALS AND METHODS

### 2.1. Iowa Mungbean Diversity Panel and Experimental Design

An Iowa Mungbean Diversity (IMD) panel with 372 genotypes was evaluated in this study. Of 372 genotypes filtered from an initial panel of 481 genotypes (Sandhu and Singh 2021) based on the Identity-by-State method (Chiteri et al. 2021). Among the nine check varieties included in the study, four elite lines (AVMU 0001, AVMU 0201, AVMU 8501, and AVMU 9701) were sourced from the World Vegetable Center, while the remaining five were commercial cultivars. The IMDP panel was tested in the experimental fields of Iowa State University’s Agricultural Engineering and Agronomy (AEA) in Boone, Iowa (coordinates: 42°00’58.5"N, 93°46’16.7"W). In 2022, the initial panel with 481 genotypes was planted on June 1; in 2023, the IMD panel with 372 genotypes was planted on June 2 at the Burkey and Bruner farms. This study used a Randomized Complete Block Design (RCBD) with two blocks in each location. Each genotype was planted in 5-foot three-row plots with 100 plants in each plot per genotype. A spacing of 2 inches between plants and 15 inches between rows was utilized. Standardized agronomic practices were implemented during this experiment, including a rain-based irrigation system.

### 2.2. Image Acquisition

All genotypes were harvested at the R6 stage when 85-90% of the pods had matured on the plants in the plot. The clusters at the top three nodes of the plants in the middle row of three-row plots with matured pods were cut using scissors and bagged in a nylon mesh bag along with the plot identifier tags and stored in the lab for image acquisition and manual data collection. An Epson Expression 11000 XL photographic scanner was used to capture RGB images of 20 randomly selected pods from each genotype with 300 dots per inch (dpi) resolution. The randomly selected 20 pods were arranged as shown in Figure 1f on a transparent plastic tray, 10 pods each to the right and left sides, and the color calibration chart in the lower center of the tray, and the plot identifier label with trial name, plot ID, and genotype ID in the top center facing inward position. Image resolution was 4299×3035 pixels, saved as 24-bit JPEGs. These components were consistently placed at the top center to enable deterministic cropping. A uniform blue background was used to aid segmentation, though lighting artifacts from the glass slide introduced complexity.

**Figure 1:**
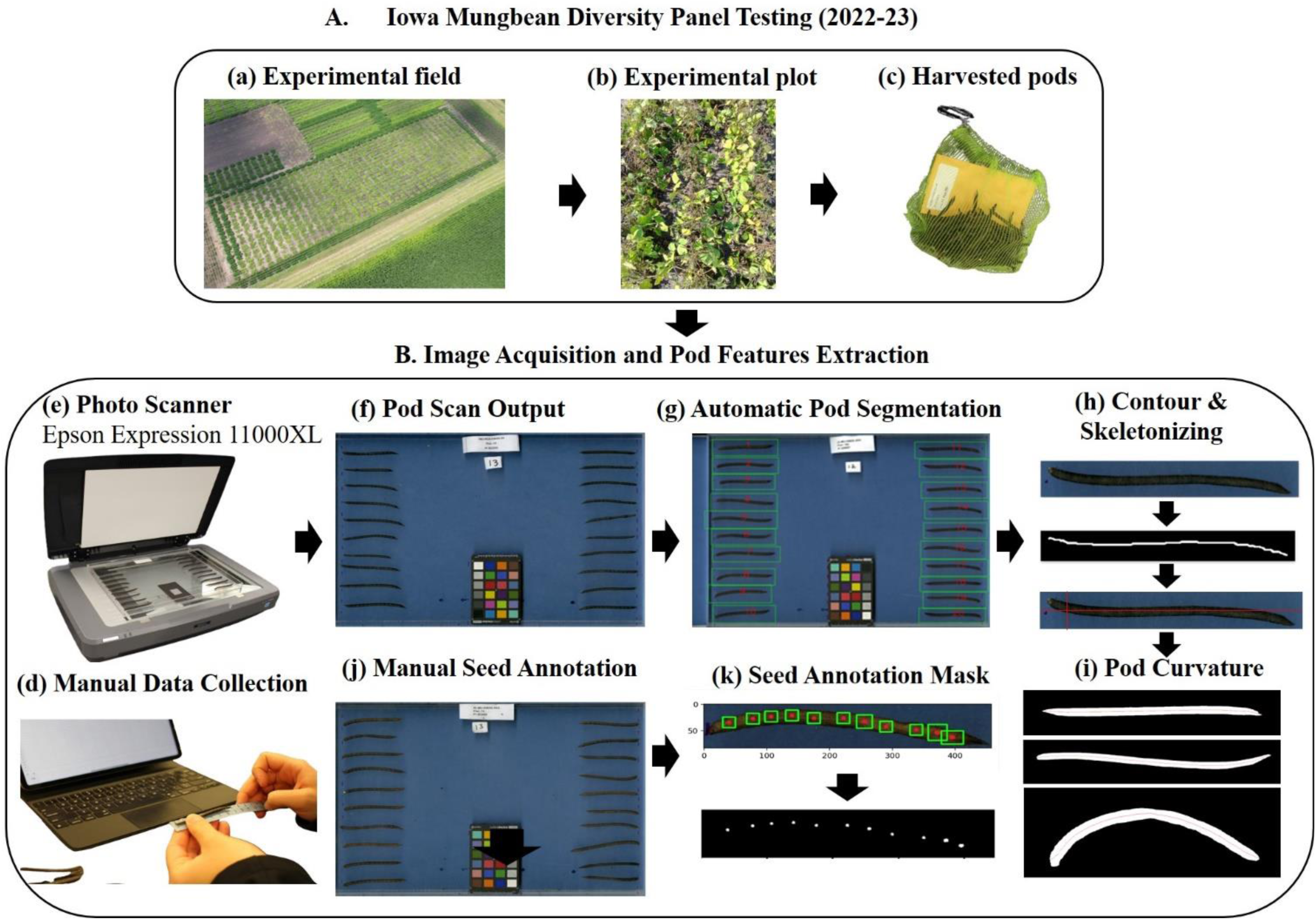
Flowchart showing the workflow for image-based mungbean pod trait extraction in the Iowa Mungbean Diversity Panel Testing (2022–23). (A) Experimental field, matured plots, and harvested pods. (B) Image acquisition and feature extraction pipeline.

### 2.3. Manual data collection and seed annotation

Pod length and seed per pod data were manually recorded to test the model accuracy. For pod curvature, the categories: (1) least curved, (2) slightly curved, and (3) curved, were utilized for manual phenotyping. Pod length was measured from the pedicle-pod joint to the tip of the pod beak using flexible scales in centimeters, using the harvested pods from two environments in 2023. The number of seeds per pod was counted without splitting the pods harvested from the 2022 and 2023 experimental plots. To train the model for extracting the number of seeds per pod, 1,678 images from each environment in 2022 and 2023 were annotated from a total of 2,418 scanned images, while images from the third location were utilized as the testing set. The scanned images were opened with Microsoft Paint, Windows 10, and the left side ten pods were annotated with red dots on the pod, based on the seed position. The data was recorded in an Excel sheet for further analysis.

### 2.4. Image analysis and Trait extraction

We developed an automated image analysis pipeline to extract morphological and seed per pod traits from scanned images containing 20 mungbean pods. The workflow comprises: (1) OCR-based metadata extraction, (2) pod detection and cropping, (3) background removal, (4) trait extraction (length, width, curvature), and (5) seed count estimation. Figure 1 illustrates the full pipeline.

#### 2.4.1. Metadata Tag Identification

Tags were consistently located in the top-middle region for pod images and the top-middle of each Petri plate. We binarized the cropped region and identified the largest connected component as the tag. OCR (Tesseract v5.3.0) was used with a restricted character set to match expected formats (e.g., Plot:1234, PI56789). Outputs were validated with regular expressions, and cropped tags were saved for traceability

#### 2.4.2. Pod Detection, Cropping, and Background Removal

Pod segmentation was achieved using a U-Net model with a ResNet34 encoder, trained using annotated data via LabelMe. The 20 largest components were sorted from bottom-left to top-right to standardize indexing. Bounding boxes were extracted to crop individual pods (Figure 1g). Each cropped pod was further processed using a DeepLabV3 model (ResNet34 backbone) for pod-background segmentation. The largest connected component was retained and refined with morphological operations to eliminate edge noise. We produced three outputs per pod: a clean binary mask, a visual overlay, and a trait-ready mask (Shen et al. 2020; Wang et al. 2022; Dhiyanesh et al. 2025; Zhou et al. 2025).

#### 2.4.3. Pod trait Extraction: Length, Width, Curvature, and seed per pod

Morphological features were extracted as follows: a) Calibration and pixel-to-physical conversion: A calibration standard (ruler or calibration card) was included in each scan to establish the pixel-to-centimeter conversion factor. The conversion factor of 1 px = 0.00847 cm was determined by measuring known distances on the calibration standard across multiple scans (n = 50) and calculating the average ratio. This corresponds to the theoretical resolution of 300 DPI (dots per inch) used by the Epson Expression scanner (1 inch = 2.54 cm; 300 pixels/inch = 0.00847 cm/pixel). The calibration was validated by measuring objects of known dimensions, achieving a measurement accuracy of ± 0.1 mm. b) Skeletonization: We inverted the binary mask and traced the midline by computing the midpoint between the topmost and bottommost contours for each x-coordinate. c) Length: The midline’s pixel length was converted to centimeters using the validated conversion factor. d) Width: 100 midline points were sampled; width was estimated as twice the shortest Euclidean distance from each midline point to the outer contour. We computed mean, max, and median widths. e) Curvature (Straightness Index): A regression line fits midline coordinates; RMSE was used as a curvature metric. f) Seed per pod estimation: A U-Net (ResNet34 encoder) was trained to segment seeds. To address noise and variability, we used an ensemble of 5 checkpoints (epochs 5– 9), averaging their outputs. Thresholded segmentations were used to count connected components, representing seeds. Ensemble averaging improved robustness, though some failure cases remain.

#### 2.4.4. Implementation and Reproducibility

The pipeline was implemented in Python 3.10 with PyTorch 1.13. Segmentation models were accessed via the segmentation models from PyTorch package. Preprocessing was handled by OpenCV and Albumentations. Outputs are saved as CSV files. All models ran on NVIDIA GPUs.

### 2.5. Descriptive statistical analysis of pod traits

The data from twenty random pods per genotype were averaged for each genotype prior to modeling. Trait distributions were inspected for normality, and Box-Cox transformations were applied where necessary. Data points that deviated from the mean by more than ±3 standard deviations were removed before analysis. Pearson correlation coefficients were calculated among pod morphological traits using mean genotype values, and visualized with the *corrplot* package, and utilized to calculate the best linear unbiased estimations (BLUEs) using the “inti” package (Lozano-Isla 2020) available in R software, using the following linear mixed model to analyze the multi-environmental data,

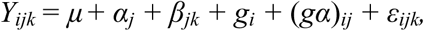

Where Y_ijk_ is the phenotypic value of the i_th_ genotype in the j_th_ environment and k_th_ block, *µ* is the overall mean, *α_j_* is the fixed effect of the j^th^ environment, 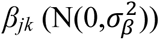 is the random effect of the k^th^ block in the j^th^ environment, *g_i_* is the fixed effect of the i^th^ genotype, 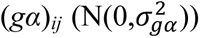 is the random effect of the i^th^ genotype in the j^th^ environment, and 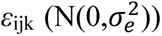 is the random error. Broad-sense heritability of these traits was estimated using the "inti" package (Lozano-Isla 2021), treating genotype as a random effect in the model described above. The calculation of broad-sense heritability was based on the equation (Cullis et al. 2006) below.

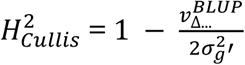

Where 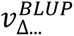 represents the mean variance of a difference between two BLUPs for the genotypic effect, and 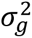 denotes the genotypic variance. We performed all the analysis using the open-source R (R Core Team 2021).

### 2.6. Linkage disequilibrium (LD) analysis and Population structure analysis

Genome-wide LD was assessed in TASSEL v5.0 (Trait Analysis by aSSociation, Evolution, and Linkage) software (Bradbury et al. 2007) using the filtered genotype file with 13414 SNPs in a HapMap format. The squared allele-frequency correlation coefficient (*r*^2^) was calculated to estimate the pairwise LD between the single-nucleotide polymorphisms (SNPs). A sliding window of 50 SNPs and a minor allele frequency (MAF) threshold of 0.05 were set to reduce bias in LD estimation. We adapted a non-linear model from Hill and Weir (1988), and implemented by Remington et al. (2001), was fitted to the data to calculate the LD decay. The model parameters, including the recombination rate (rho), were estimated using non-linear least squares (NLS) regression. The LD values were adjusted based on their distances along the chromosome. The half-decay distance, which is the point where LD decreases to half of its maximum value, was estimated. The results were then visualized by plotting the observed and adjusted r² values, along with the calculated half-decay distance using the R package.

To assess the population structure of the IMD panel, a Bayesian statistic-based clustering method was implemented in STRUCTURE v2.3.4 (Pritchard et al. 2000). A Total of 13414 SNP markers were used for this analysis. The admixture model with correlated allele frequencies was applied, assuming a range of possible subpopulations (K) from 2 to 10. Five runs were performed independently for each K with a burn-in period of 10,000 iterations, followed by 100,000 Markov Chain Monte Carlo (MCMC) replications. The optimal K value was determined using the ΔK method (Evanno et al. 2005) implemented in STRUCTURE HARVESTER (Earl and vonHoldt 2012). The CLUMPAK method (Kopelman et al. 2015) was utilized to account for run-to-run variation to visualize the results.

### 2.7. Phenotypic Principal Component Analysis

Principal component analysis (PCA) was conducted to examine the phenotypic structure and relationships among various pod morphological traits. Seven pod traits (image and manual data) were analyzed using the BLUEs from 372 genotypes to ensure consistency for subsequent genetic association analyses. Before performing PCA, all trait values were standardized (mean = 0, standard deviation = 1) to ensure each trait was given equal weight, regardless of its measurement scale. Based on the column means, the missing values were imputed. PCA was executed using singular value decomposition to extract the principal components, with the number of components retained based on eigenvalues greater than 1 and the cumulative variance explained. The contribution of each trait to the principal components was assessed based on the component loading calculations. The analysis was carried out using the scikit-learn package in Python (version 1.3.0) (Pedregosa et al., 2011). The variance explained by each component, as well as the cumulative variance, was calculated to evaluate the dimensionality of phenotypic variation. A biplot visualization was created to display both genotype scores and trait loadings within the principal component space.

### 2.8. Genome-Wide Association Analysis

Association mapping was conducted for pod morphological traits, including pod length (cm), width (mm), curvature, and the number of seeds per pod. The Best Linear Unbiased Estimations (BLUEs) calculated using both manually collected and image-based data were utilized for GWAS. In this study, we compared a single-locus model: MLM (Zhang et al. 2010) and three multi-locus models: farmCPU (Liu et al. 2016), Bayesian Information and Linkage-disequilibrium Iteratively Nested Keyway (BLINK) (Huang et al. 2019) using the GAPIT (version 3) (Wang and Zhang 2021) package and Selection of Variables with Embedded screening using Bayesian methods (SVEN) (Li et al. 2023) for association studies. The initial IMD panel was genotyped using genotype-by-sequencing, resulting in 26550 single-nucleotide polymorphisms (SNPs) (Sandhu and Singh 2021) for 500 genotypes, and the shortlisted panel with 372 genotypes selected for this study is left with 19658 SNPs. For this study, a total of 13414 SNPs remained for analysis after filtering out sites with less than 15% missing data and 0.05 minor allele frequency from the 19658 SNP markers. These SNPs were utilized for calculating the kinship matrix (K matrix) and population structure (first three principal components(PC)) in Trait analysis by ASSociation, Evolution, and Linkage (TASSEL) (Bradbury et al. 2007) to fit as covariates in the model in GAPIT for GWAS analysis to address the confounding effects due to population structure and relatedness. The GWAS results, except SVEN, the other three models were visualized using Manhattan plots, utilizing the ggplot2, ggrepel, and ggpubr packages. Q-Q plots were created with the qqman package (Turner 2014). A genome-wide significance threshold of −log_10_ (0.05/*n*) was set, where *n* is the number of comparisons, following the Bonferroni correction to control for multiple comparisons.

### 2.9. Candidate gene identification and comparative mapping

Two common SNPs associated with pod morphological traits were identified for downstream analysis. Candidate gene searches were performed using the Legume Information System (LIS) JBrowse tools (https://www.legumeinfo.org/) to identify genes within the LD block around the significant SNP region. This analysis was based on the average linkage disequilibrium (LD) decay observed in the population. The identified protein sequences were compared using BLASTP (Wheeler and Bhagwat 2007) with the protein databases of soybean (*Glycine max*), cowpea(*Vigna unguiculate*(L)), and common bean (*Phaseolus vulgaris*) available in the Legume Information System and with *Arabidopsis thaliana* in The Arabidopsis Information Resource (TAIR-https://www.arabidopsis.org/tools/blast/) using an E-value cutoff of 1e-20. The identified matches from the *Arabidopsis* were evaluated for the functional descriptions related to these traits. A chord diagram to visualize synteny between mungbean, soybean, cowpea, and common bean proteins was generated on the SequenceServer (version 3.1.2) using BLASTP 2.15.0 (Priyam et al. 2019) in LIS (https://sequenceserver.legumeinfo.org/). Additional comparative mapping tools available on the LIS: ZZBrowse, JBrowse, Genome Context Viewer, Gene Family Search (Dash et al. 2016; Redsun et al. 2022) were used for comparative genomic analysis.

### 2.10. Genomic Prediction

To evaluate the accuracy of genomic prediction (GP) for pod morphological traits, a ridge regression best linear unbiased prediction (rrBLUP) model was employed using the rrBLUP package (Endelman 2011). Genomic predictions were conducted using a marker matrix derived from 13414 SNP markers and phenotypic values collected through both manual and image-based phenotyping methods. For each trait, we performed ten-fold cross-validation (CV), which balances bias and variance in prediction accuracy estimates and maintains adequate training set size for our population of 372 genotypes. The phenotype values (trait BLUPs) for each genotype were generated from across environments were randomly divided into ten equal-sized subsets, ensuring balanced representation across the diversity panel. In each iteration, nine subsets (90%) were used to train the model, while the remaining subset (10%) was used to test prediction accuracy. This process was repeated ten times so that each sample was used once for validation. Genomic prediction accuracy (GPA) was evaluated using the Pearson correlation coefficient, coefficient of determination (R^2^), mean squared error (MSE), and prediction bias between observed and predicted phenotypes. In each CV fold, the Pearson correlation coefficient was calculated between the genomic estimated breeding values (GEBVs) predicted by the rrBLUP model and the observed trait BLUPs for each genotype in the test set. In each fold, a minor allele frequency (MAF) filter (≥ 0.05) was applied to remove rare SNPs that could cause overfitting, and markers with zero variance within the training set for each fold were excluded to maintain numerical stability. Performance metrics were calculated per fold and averaged across ten folds, and standard errors were computed to assess prediction reliability. To enable robust comparison between image-based and manual phenotyping approaches, identical CV partitions were used within each trait. A selection coincidence analysis was also conducted to assess agreement between phenotypic and genomic prediction. For each trait, the top ten genotypes based on both phenotypic BLUPs and GEBVs were identified, and the selection coincidence index was calculated based on the proportion of the genotypes common to both top-10 lists (coincidence = number of common genotypes/10).

## 3. RESULTS

### 3.1. Image analysis and trait extraction accuracy

Our machine learning-based analysis model showed high accuracy in quantifying the morphological traits of mungbean pods and seeds. Pod length measurements (r = 0.96, p < 0.001) and seeds per pod counts (r = 0.963, p < 0.001) showed exceptionally high agreement between image-based and manual measurement methods. (Figure 2). This demonstrates the model accuracy and confirms the reliability of the automated phenotyping approach. Since manual measurements for pod width were impractical, we relied solely on the model for this data in our further analysis.

**Figure 2:**
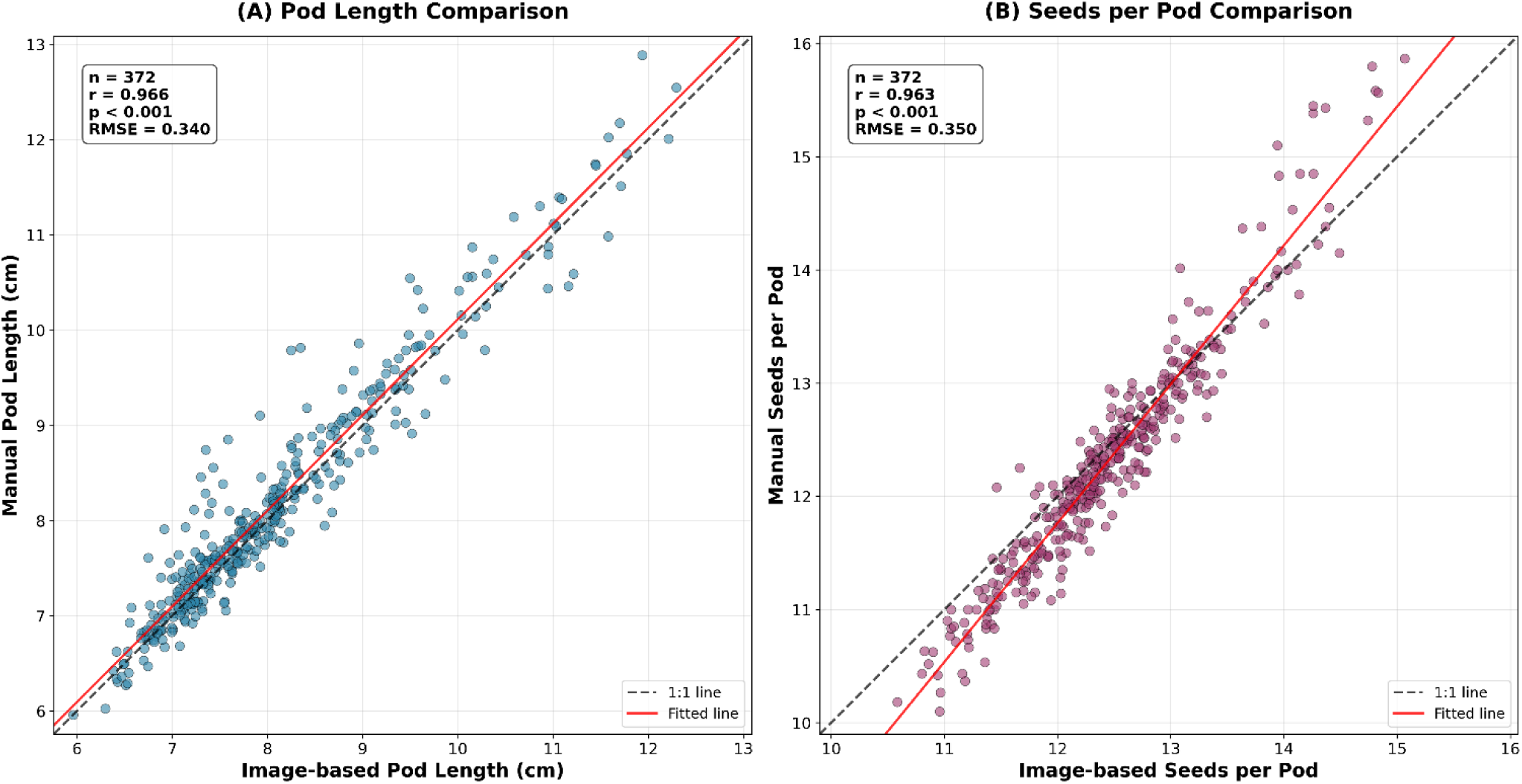
Validation of image-based pod phenotyping accuracy. Correlation between image-based and manual measurements for (A) pod length (n = 372, r = 0.966, RMSE = 0.34 cm) and (B) seeds per pod (n = 372, r = 0.963, RMSE = 0.35 seeds) across mungbean genotypes from multiple environments. The 1:1 line (black dashed) represents perfect agreement, and the fitted regression line (red solid) shows the actual relationship. Both traits demonstrate excellent accuracy for high-throughput phenotyping applications.

### 3.2. Phenotypic variation and trait relationships

A comprehensive phenotypic evaluation of pod and seed traits, based on both image analysis and manual measurements, revealed significant variation among the IMD panel accessions for these traits (see Table 1, Figure 3). For pod length, image-based measurements ranged from 5.96 to 12.29 cm, with a mean of 8.09 cm and a standard deviation of 1.21 cm, and a high broad-sense heritability (H^2^ = 0.91). Manual measurements of pod length showed similar results, with a mean of 8.19 cm, a standard deviation of 1.26 cm, and a range of 5.96 to 12.89 cm, but slightly lower heritability (H^2^ = 0.81). Pod width, measured only through image analysis, ranged from 2.80 to 4.94 mm, with a mean of 3.79 mm and a standard deviation of 0.37 mm, also exhibiting high heritability (H^2^ = 0.91). Pod curvature, assessed via image analysis, showed considerable variation, with a range of 0.05 to 0.24, a mean of 0.09, and a coefficient of variation (CV) of 0.35. It demonstrated strong genetic control (H^2^ = 0.83). Manual assessments of pod curvature reported lower absolute values but similar trends, with a mean of 1.98 and heritability of H^2^ = 0.55. Seed count per pod exhibited moderate variation, with manual measurements showing a mean of 12.45 seeds (range: 10.58 to 15.07 and CV = 0.06) and image measurements displaying a mean of 12.31 seeds (range: 10.10 to 15.87 and CV = 0.08). Both methods demonstrated moderate heritability (manual H^2^ = 0.75; image H^2^ = 0.74). Among all traits evaluated, pod length displayed the largest standard deviation, followed by seeds per pod, pod width, and pod curvature. Overall, genotype variance consistently surpassed environmental and GxE components, highlighting the significant genetic influence on phenotypic expression (Table 1).

**Table 1:**
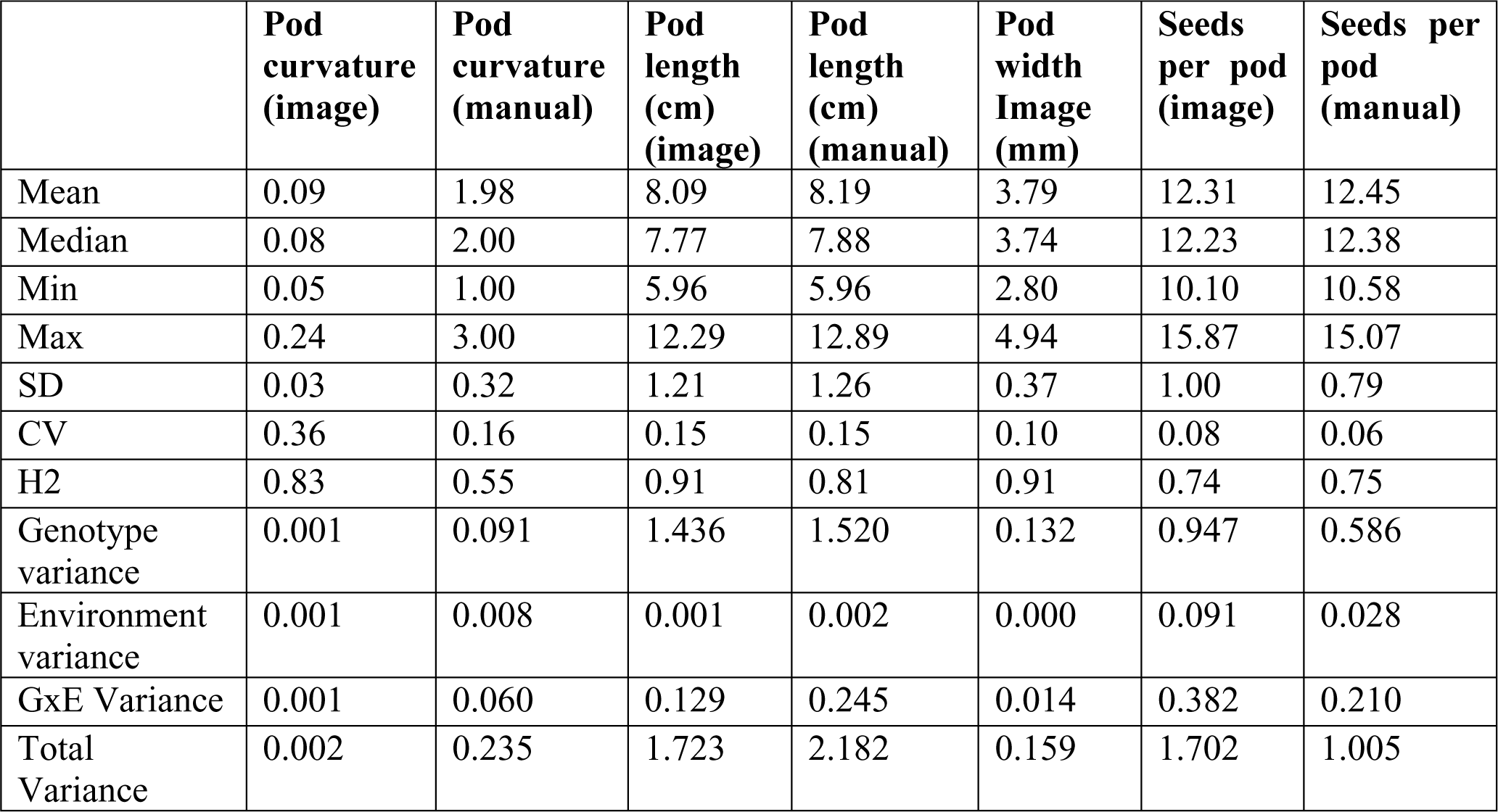
Descriptive statistics, coefficients of variation (CV), and broad-sense heritability (H^2^) of the pod morphological traits derived from image-based phenotyping.

|  | **Pod curvature (image)** | **Pod curvature (manual)** | **Pod length (cm) (image)** | **Pod length (cm) (manual)** | **Pod width Image (mm)** | **Seeds per pod (image)** | **Seeds per pod (manual)** |
| --- | --- | --- | --- | --- | --- | --- | --- |
| Mean | 0.09 | 1.98 | 8.09 | 8.19 | 3.79 | 12.31 | 12.45 |
| Median | 0.08 | 2.00 | 7.77 | 7.88 | 3.74 | 12.23 | 12.38 |
| Min | 0.05 | 1.00 | 5.96 | 5.96 | 2.80 | 10.10 | 10.58 |
| Max | 0.24 | 3.00 | 12.29 | 12.89 | 4.94 | 15.87 | 15.07 |
| SD | 0.03 | 0.32 | 1.21 | 1.26 | 0.37 | 1.00 | 0.79 |
| CV | 0.36 | 0.16 | 0.15 | 0.15 | 0.10 | 0.08 | 0.06 |
| H2 | 0.83 | 0.55 | 0.91 | 0.81 | 0.91 | 0.74 | 0.75 |
| Genotype variance | 0.001 | 0.091 | 1.436 | 1.520 | 0.132 | 0.947 | 0.586 |
| Environment variance | 0.001 | 0.008 | 0.001 | 0.002 | 0.000 | 0.091 | 0.028 |
| GxE Variance | 0.001 | 0.060 | 0.129 | 0.245 | 0.014 | 0.382 | 0.210 |
| Total Variance | 0.002 | 0.235 | 1.723 | 2.182 | 0.159 | 1.702 | 1.005 |

Correlation analysis (Figure 3) revealed strong positive relationships between pod dimensional traits, highlighting their coordinated development. Among pod dimensional traits, pod length (both manual and image-based) and width were strongly correlated (r = 0.77 −0.80), and pod curvature (image-based) was moderately correlated with width (r = 0.61) and pod length (image-based and manual) (r = ∼ 0.73) but manual based curvature showed lower correlation (r = 0.45) with width. In contrast, seed per pod (both manual and image-based) showed weak correlations with pod curvature (both manual (r = 0.27 - 0.35 and image-based (r = 0.27 - 0.36) and pod width (r = 0.28 - 0.32), suggesting a distinct and partially independent genetic architecture but a moderate correlation with pod length (both manual and image-based r = 0.56 - 0.63). All correlation coefficients were statistically significant at p < 0.001 (Figure 3).

**Figure 3:**
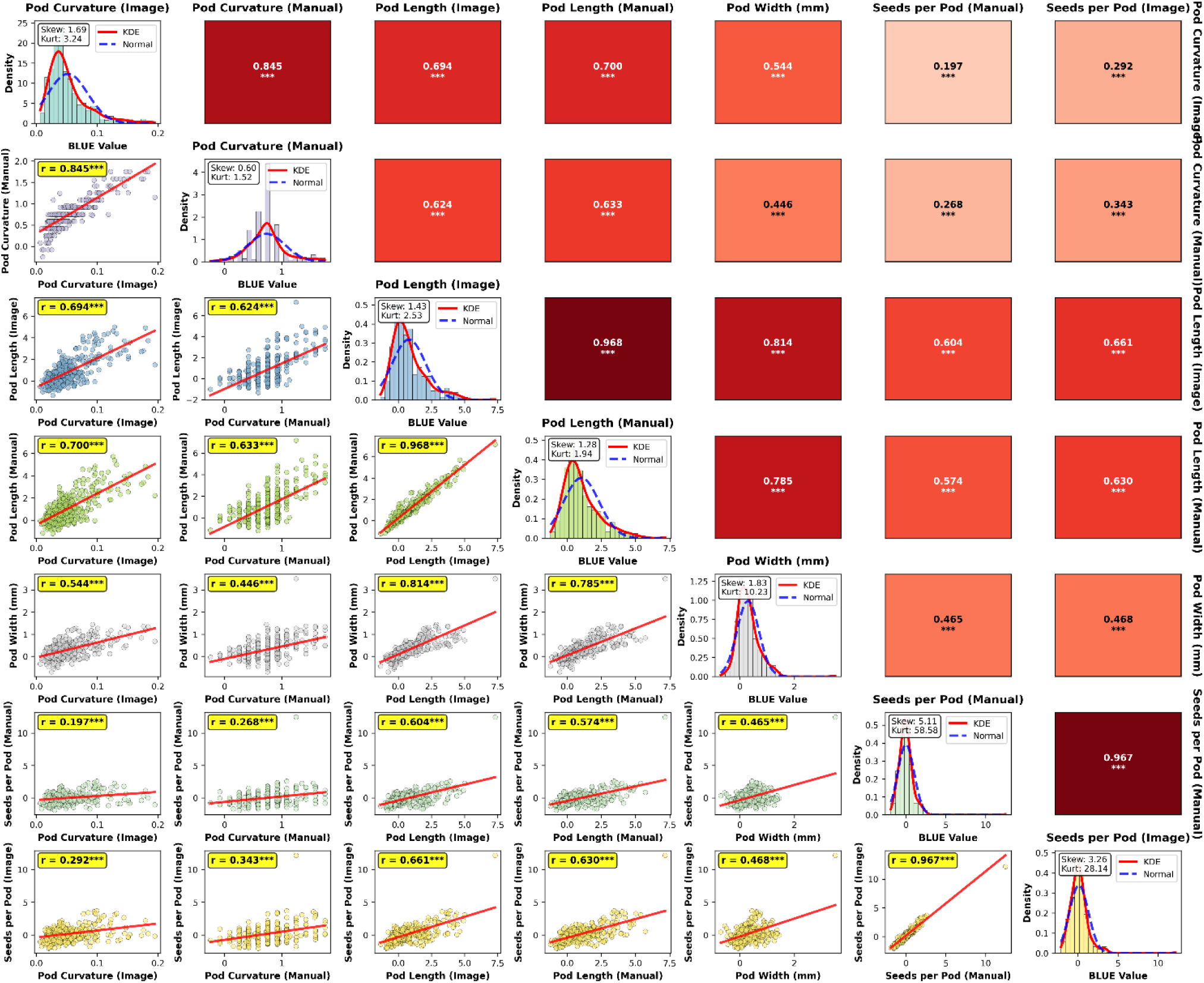
Correlation matrix and phenotypic distributions of pod and seed traits. Heatmap showing pairwise Pearson correlation coefficients among eight pod and seed traits. Stronger correlations are indicated by darker shades of red. Histograms showing the distribution of individual traits.

### 3.3. Linkage disequilibrium analysis and population structure analysis

The linkage disequilibrium (LD) was estimated between 13414 SNP pairs mapped to 11 chromosomes of mungbean. Figure 4A illustrates the LD decay plot. At approximately 266 kb of physical distance, the LD dropped below the threshold of r² = 0.2. Population structure analysis conducted with the STRUCTURE software identified four distinct subpopulations (K=4) within the IMD panel (Figures 4B and 4C). Among these subpopulations, population one (blue dots) comprises approximately 50% of the genotypes, followed by population two (orange dots) at about 30%, population three (green dots) at around 17%, and population four (red dots) at 4%. Principal Component Analysis (PCA) further confirmed the existence of structured genetic variation within the panel (Figure 4D). The first two principal components (PC1 and PC2) accounted for 10.4% and 5.0% of the total variance, respectively, and showed a partial clustering that corresponds with the subpopulations defined by STRUCTURE.

**Figure 4:**
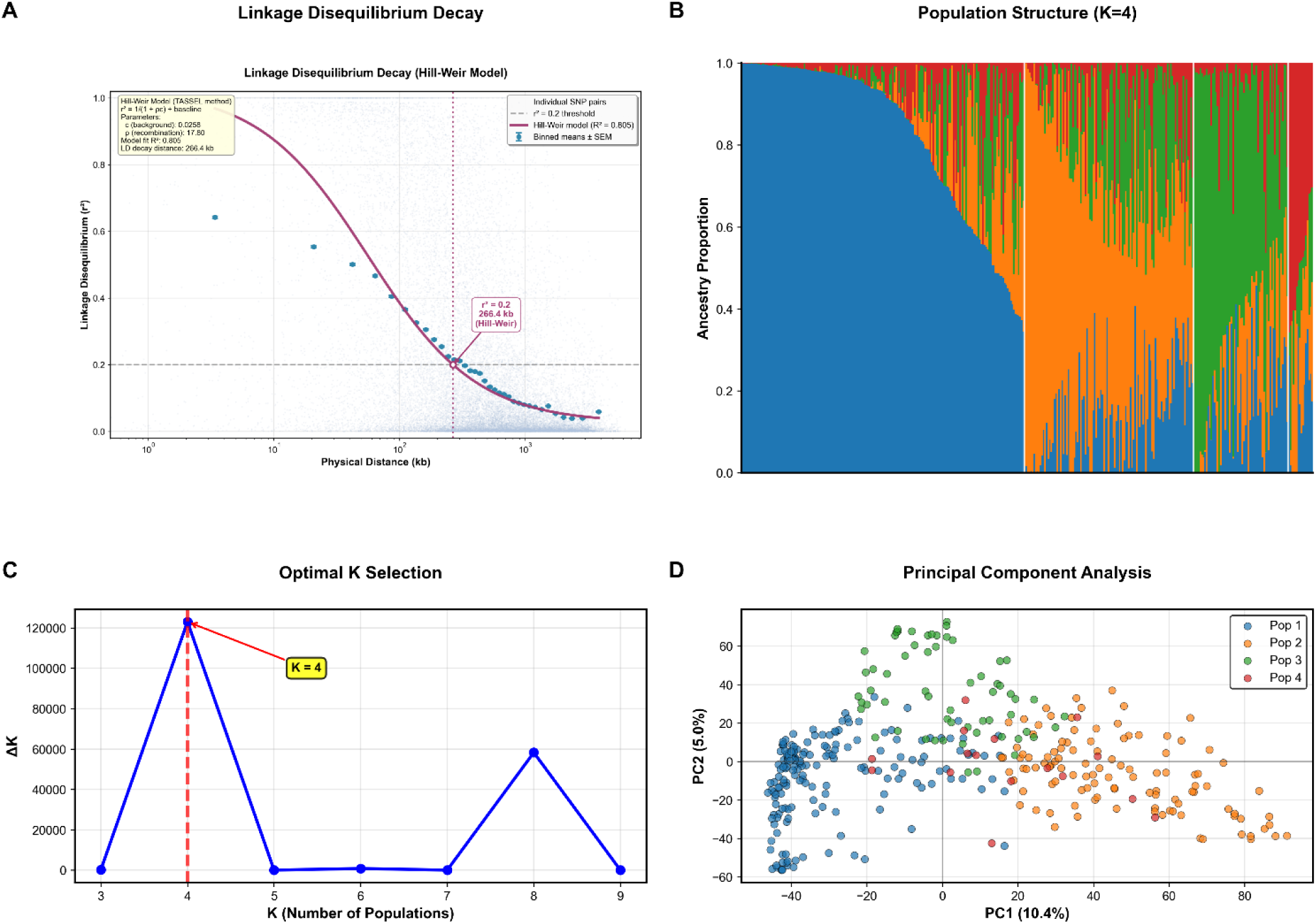
LD analysis and population structure of the Iowa mungbean diversity panel. A) LD decay curve showing r^2^ values against physical distance. B) Bar plot of population structure at K = 4, with individuals partitioned into four clusters. C) Delta K plot supporting K = 4 as the optimal number of subpopulations. D) Principal Component Analysis (PCA) plot showing the percentage of variation explained by the first two principal components.

### 3.4. Phenotypic structure and principal component analysis

Principal component analysis revealed distinct patterns of phenotypic variation among the seven pod morphological traits. The first two principal components explained 85.2% of the total phenotypic variance (PC1: 65.8%, PC2: 19.4%), indicating that the majority of morphological variations could be effectively captured in a two-dimensional space (Figure 5). PC1 was primarily associated with overall pod size characteristics, with the strongest positive loadings from pod length measurements (image-based: +0.445, manual: +0.441), pod curvature measurements (image-based: +0.379, manual: +0.347), and pod width (+0.358). This component represented a general pod morphology factor, with larger pods tending to be longer, wider, and more curved. PC2 was characterized by a contrast between seed per pod and pod curvature characteristics. Seeds per pod measurements showed strong positive loadings (manual: +0.614, image-based: +0.561), while pod curvature measurements exhibited negative loadings (image-based: −0.365, manual: 0.328). This component distinguishes genotypes with high seed production capacity from those with pronounced pod curvature. The third principal component explained an additional 9.6% of variance (cumulative: 94.8%), primarily capturing residual variation in pod width relative to other morphological features.

**Figure 5.**
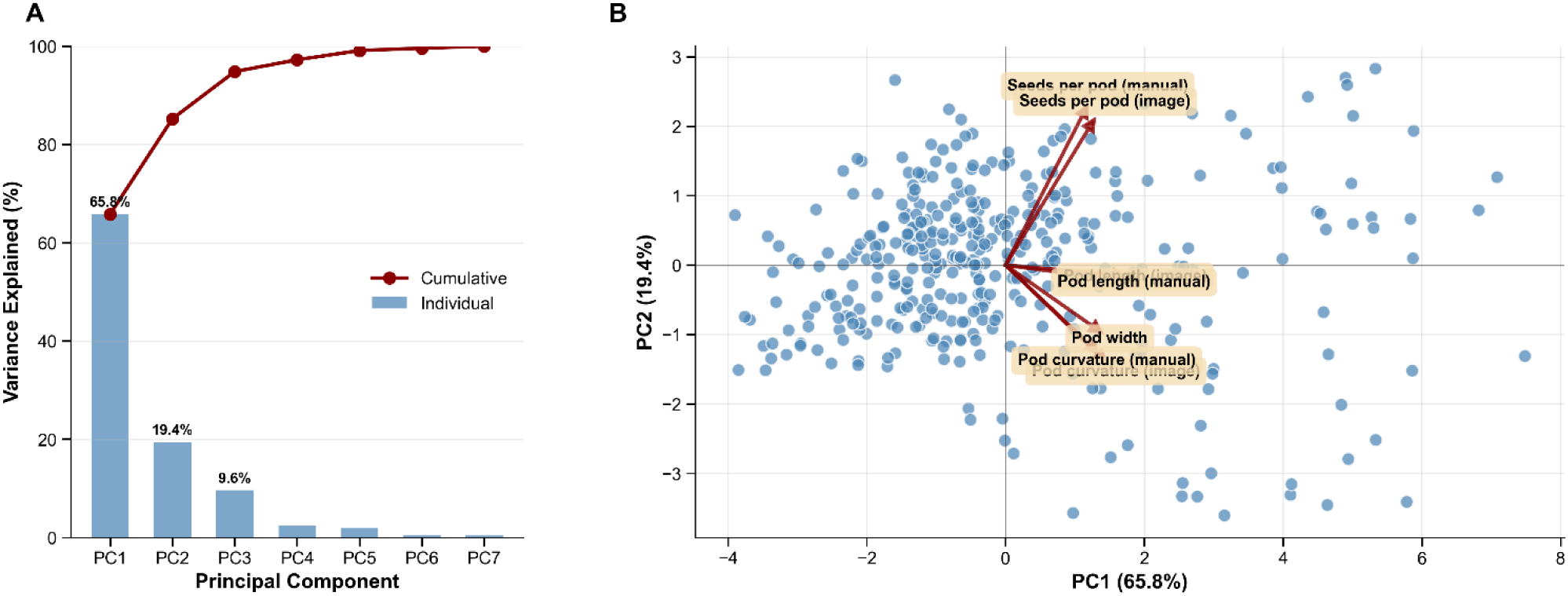
Principal component analysis based on BLUEs values from 372 genotypes across seven morphological traits. (A) Scree plot showing individual and cumulative variance explained by principal components. (B) PCA biplot displaying genotype scores (blue points) and trait loading vectors (red arrows) in PC1-PC2 space. PC1 (65.8% variance) represents overall pod morphology, while PC2 (19.4% variance) contrasts seed per pod with pod curvature traits.

This phenotypic structure analysis provided the foundation for subsequent genome-wide association studies with BLUEs, ensuring consistency between phenotypic characterization and genetic association mapping approaches (Section 3.5).

### 3.5. Genome-Wide Association Study (GWAS) results

To unravel the genetic architecture underlying the observed phenotypic variation in pod morphological traits, GWAS was conducted using both manually measured and image-analysis-based predicted datasets.

#### 3.5.1. Comparison of manual and image-based phenotyping data in GWAS

The phenotypic data on pod curvature, length, width, and seeds per pod, obtained from the image-analysis model, showed promising results from the GWAS analysis using four different models: FarmCPU, BLINK, MLM, and SVEN. These results were compared to those obtained from manual data-driven GWAS (see Figure 5 and Table S1). We identified common significant single-nucleotide polymorphisms (SNPs) for all traits, with the number of significant SNPs derived from the image-based data-driven GWAS exceeding that from the manual data-based analysis (see Figures 5A-C and Table S1). The most significant SNPs identified were located on chromosome 1, followed by chromosomes 3 and 2 (Figure 5C). For pod length, the image-based GWAS identified 14 significant SNPs, with four SNPs colocalized between FarmCPU and BLINK, and one SNP common to both FarmCPU and MLM methods. In contrast, the manual data-driven GWAS identified a total of eight SNPs for pod length. For seed per pod, the image-based GWAS yielded ten significant SNPs, with one SNP common between FarmCPU and BLINK, whereas the manual approach also identified ten significant SNPs, with no common SNPs across the three methods. Similarly, the analysis of image-based data for pod curvature revealed 13 significant SNPs, including one SNP identified across all three methods. In contrast, the manual data for pod curvature resulted in eight SNPs, with one common SNP found between FarmCPU and BLINK.

**Figure 5:**
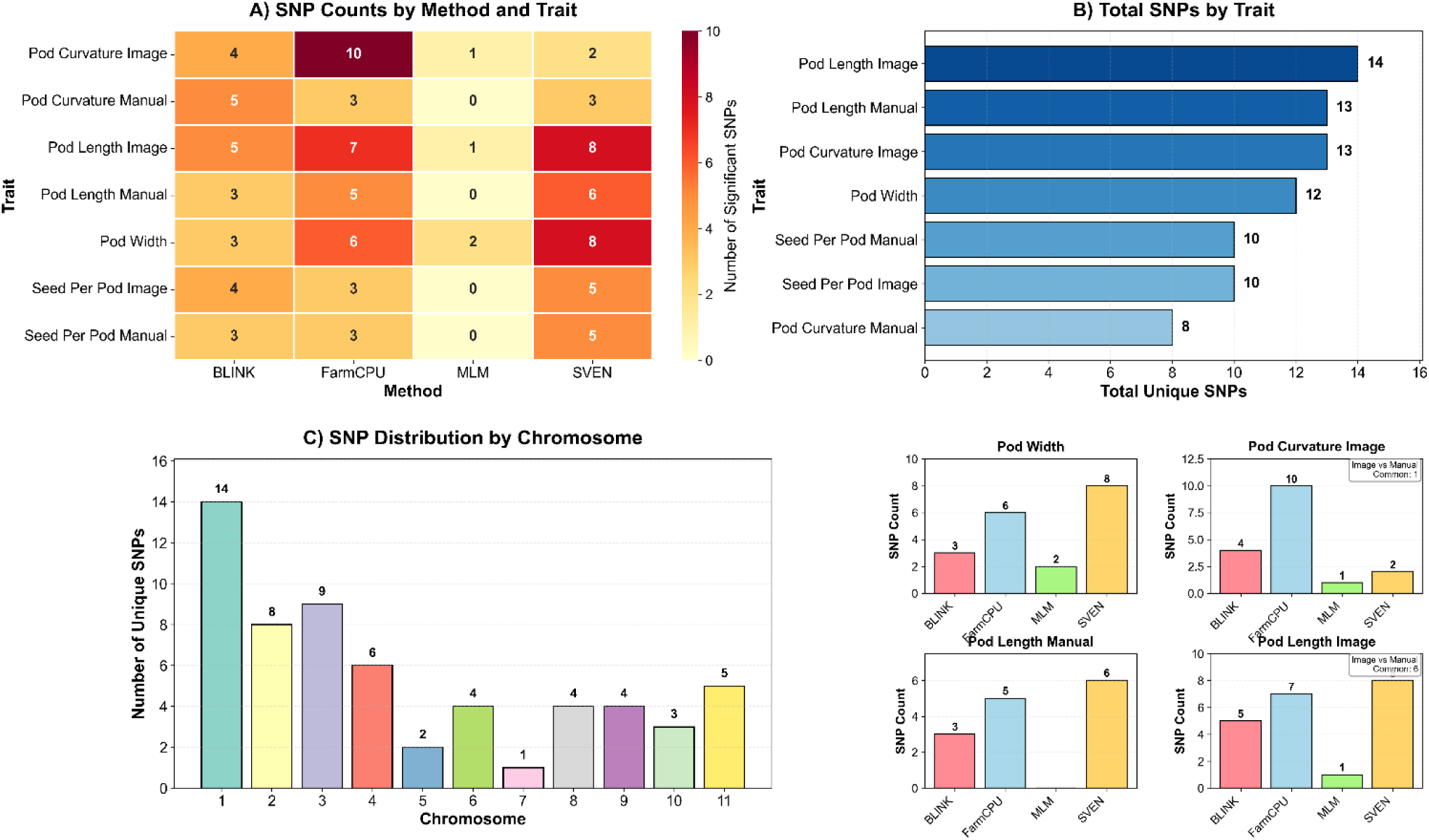
Comprehensive comparison of four GWAS methods (BLINK, FarmCPU, MLM and SVEN) for SNP discovery across pod morphology traits in the IMD panel. (A) SNP counts by method and trait showing FarmCPU’s superior performance. (B) Total number of SNPs for each trait, including four methods. (C) Chromosomal distribution of significant associations (D). Bar plots showing the Total SNPs among the GWAS methods for four traits and the overlap between both the image-based and manual methods of phenotyping.

#### 3.5.2. Genome-wide association and candidate gene identification

In this study, we employed MLM, BLINK, FarmCPU, and SVEN methods for association mapping, which led to the identification of varying numbers of significant markers associated with pod morphological traits. Given the established agreement between manual and image-based data, along with the superior accuracy of the image-based data (see Figure 5 and Table S1), we further discuss the GWAS results that are driven by image-based analysis. Multiple SNPs were found to be co-localized across these methods, enhancing our confidence in these loci. This analysis identified a total of 45 significant SNPs for all the pod morphological traits (see Table S1). Manhattan and corresponding Q-Q plots were generated using results from the MLM, BLINK, and Farm CPU methods (Figure 6). Note that the naming of SNPs is based on their chromosome number and their position on the chromosome according to the reference genome version 6, published by Kang et al. (2014).

**Figure 6:**
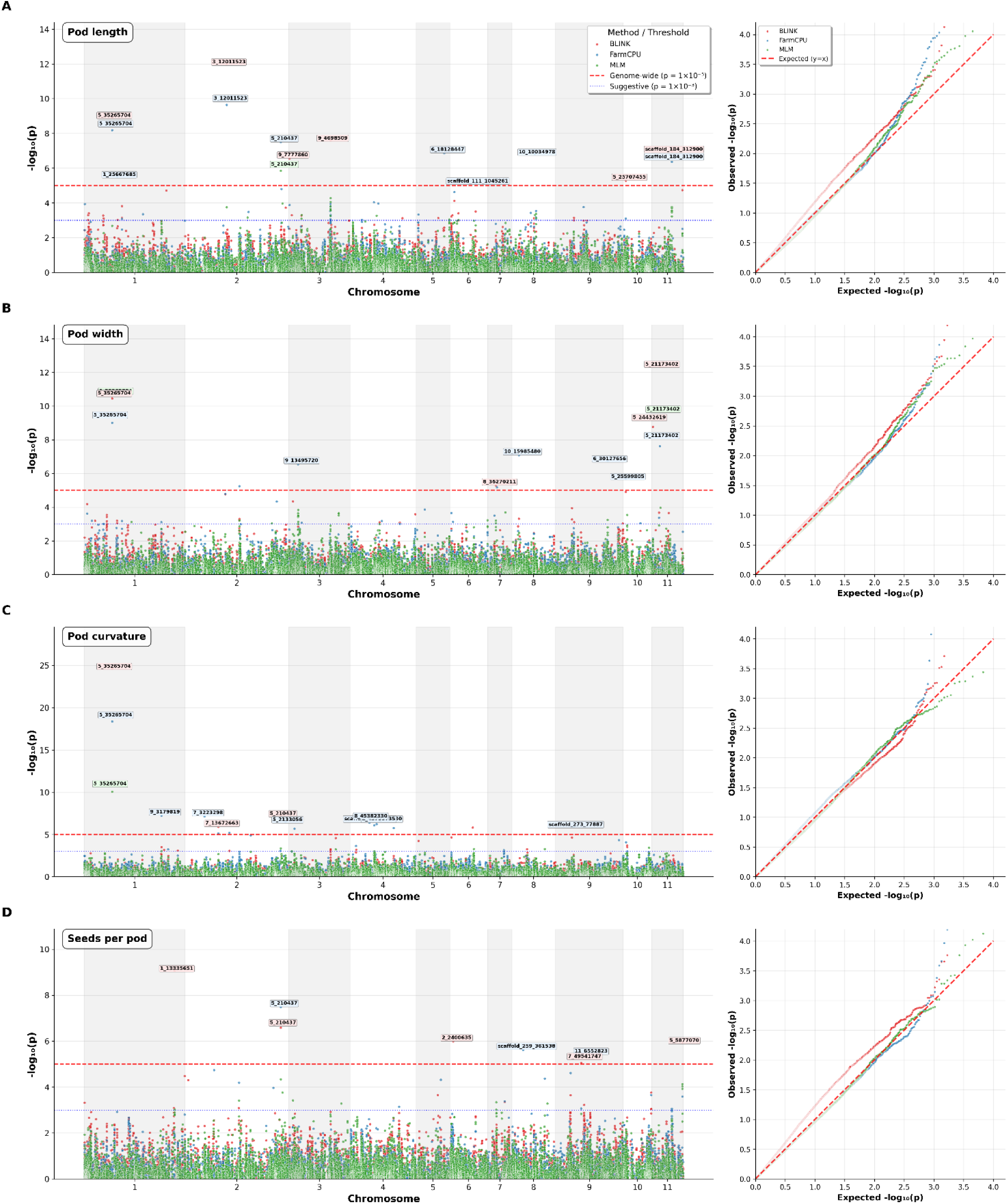
Manhattan and corresponding Q-Q plots that illustrate the results of genome-wide association studies (GWAS) based on the FarmCPU, BLINK, and MLM methods, focusing on image-based phenotypic traits like pod length, width, curvature, and seeds per pod. The red dashed line in the Manhattan plots indicates the Bonferroni genome-wide correction threshold set at α = 0.05. The red dashed line in Q-Q plots represents the expected distribution of p-values under the null hypothesis of no association. Deviations above the red line indicate SNPs with stronger associations than expected by chance.

For pod curvature, GWAS analysis employing these three models identified 13 SNPs, which explained phenotypic variation (PVE) of a range between 11.28 and 54%. Out of these 13 SNPs, four SNPs are located on chromosome 2 (5_210437 (PVE:4.7-6.15%), 7_13672663 (3.02%), 7_3223298 (2.13%), and 5_2133056 (1.58%), four on chromosome 4 (scaffold_309_152830 (0%), 8_45382330 (0%), 8_43793735 (0.68%) and 3_9992892 (0%), two on chromosome 1 (5_35265704 (2.25 – 54.51%) and 9_3179819 (0%)) and the rest harbored one SNP each on chromosome 3 (9_9820033 (4.64%)), 6 (2_15269420 (6%)) and 9 (scaffold_273_77887 (0%)). An SNP 5_35265704 on chromosome 1 was colocalized among the four models, and another SNP 5_210437 on chromosome 2 was between the BLINK and SVEN models.

For pod length, these four models identified a total of 14 SNPs associated with pod length, which explained between 9% and 53% of the total phenotypic variance. Among these, SNP 5_210437 on chromosome 2 was found to be common between the MLM, SVEN, and FarmCPU models, with 6.05%, 11.1% and 13.67% PVE, respectively. Additionally, two SNPs were identified as common between the FarmCPU and BLINK models, namely, SNP 5_35265704 on chromosome 1 showed PVE values of 1.63% and 18.17% respectively, and SNP 3_12011523 on chromosome 2 had PVE values of 0.33% and 11.94%. An SNP, scaffold_184_312900 on chromosome 11, was common among BLINK, FarmCPU, and SVEN models, and with PVE of 0.97%, 9.59% and 6.6, respectively, and another SNP, 10_10034978, is common between FarmCPU and SVEN models with PVE of 0% and 4.9% respectively. Of the remaining ten other significant SNPs, three were harbored on chromosome 3 (9_7777860 (5.87%), 9_4698509 (8.02%), and scaffold_387_2663 (0.8%), and two were on chromosome 1 (1_25667685 (0%) and 1_33591256 (37%). Out of the remaining four SNPs, SNP 5_669301(11.8%) was located on chromosome 2, SNP 7_22470394 (8.8%) on chromosome 4, SNP 6_18128447 (0%) on chromosome 5, and SNP 5_25707455 (13.6%) on chromosome 10.

For pod width, four models identified 12 SNPs across eight chromosomes. Together, these SNPs explain 27% to 72% of the phenotypic variance, with the highest contribution recorded in the SVEN model (72%), followed by the BLINK (70%) and MLM (65%) models. One SNP, 5_35265704, located on chromosome 1, was identified by all four methods and accounted for a larger portion of the phenotypic variance (1.92% to 51%). Another SNP, 5_21173402, on chromosome 11, was also identified by all four models, with a PVE range of 11.75% to 20.1%. In addition, an SNP 5_24432619 on chromosome 11 was also colocalized between BLINK and SVEN models with a PVE of around 14%. Out of eight remaining SNPS, two were located on chromosomes 3 (9_8662740 (19.1%) and 9_13495720 (4.4%). The remaining six SNPs were located on chromosome 1 (1_7036436 (9.7%)), chromosome 6 (2_14494712 (27.1%), 8 (10_15985480 (0%)), 9 (6_30127656 (0%)), 10 (5_25599805 (4.59%)), 11 (5_24432619 (0.3%)).

For the seed per pod trait, except MLM, all three other models identified ten significant SNPs, with a PVE ranging from 5.18% to 48%. Of these ten SNPs, three (1_5295485 (2%), 1_11367629 (13.7%), and 1_13335651 (10.1%)) were located on chromosome 1, and two (7_49541747 (12%) and 11_6552823 (2.65%)) were on chromosome 9. Of the remaining five, SNP 7_20491843 (4%) was on chromosome 2, SNP 2_2400635 (4.18%) is on chromosome 6, scaffold_259_361538 (0%) is on chromosome 8, and SNP 11_65528235 (2.65%) is on chromosome 9. Interestingly, an SNP 5_210437 on chromosome 2, colocalized by the BLINK, FarmCPU, and SVEN methods, explained a phenotypic variance of 2.52%, 12.3% and 15.4%, respectively.

In Table 2, we presented the top two to three SNPs for each image-based trait, along with their allelic effect, minor allelic frequency, and PVE. In addition, we extracted the genes from the reference genome using an LD block window of 532 Kb (which includes 266 Kb upstream and downstream). Based on gene ontology information, we identified potential candidate genes for each trait and provided their orthologs in *Arabidopsis*, along with their protein names and descriptions sourced from the TAIR website(Schäffer et al. 2001) (https://www.arabidopsis.org/tools/blast/).

**Table 2:**
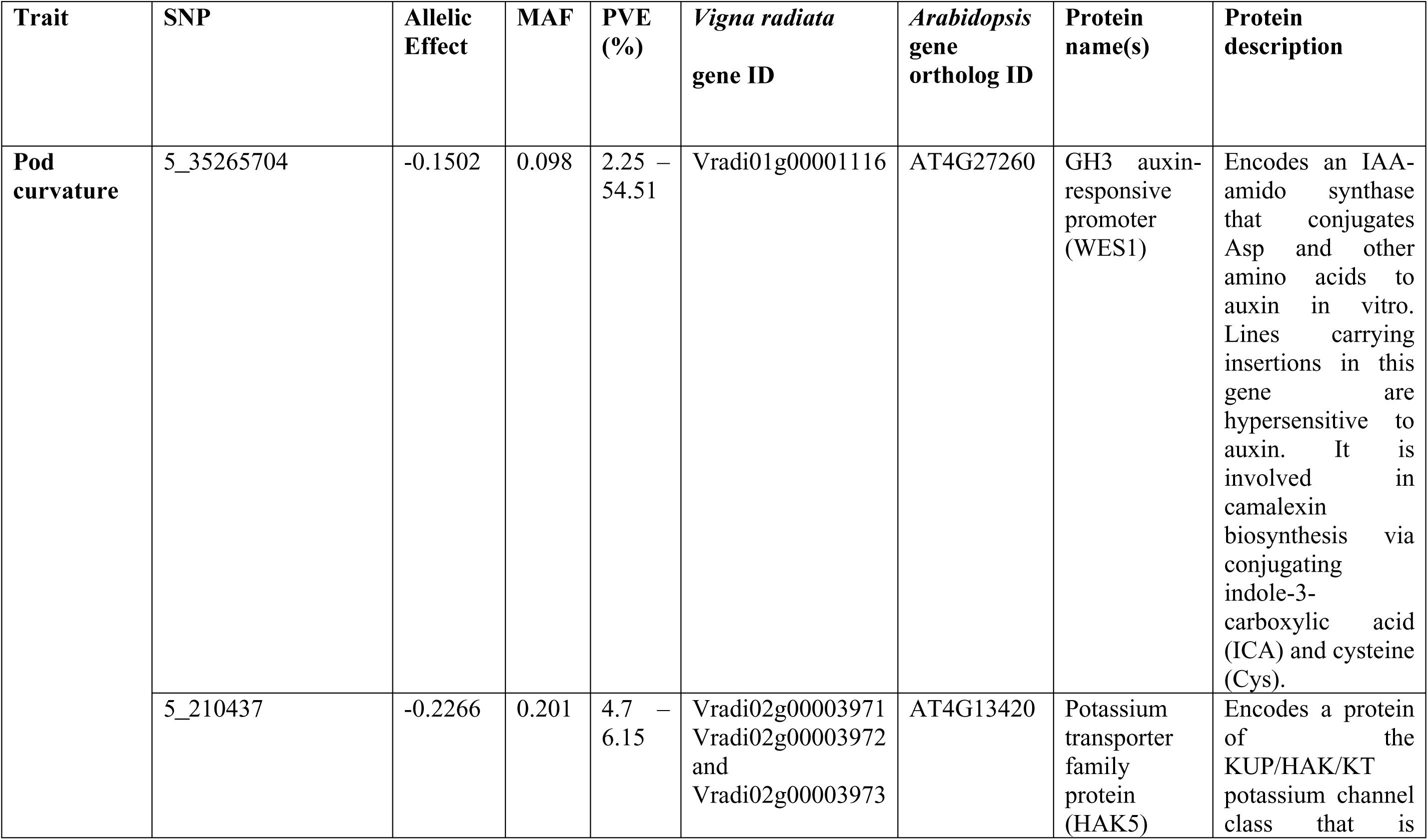
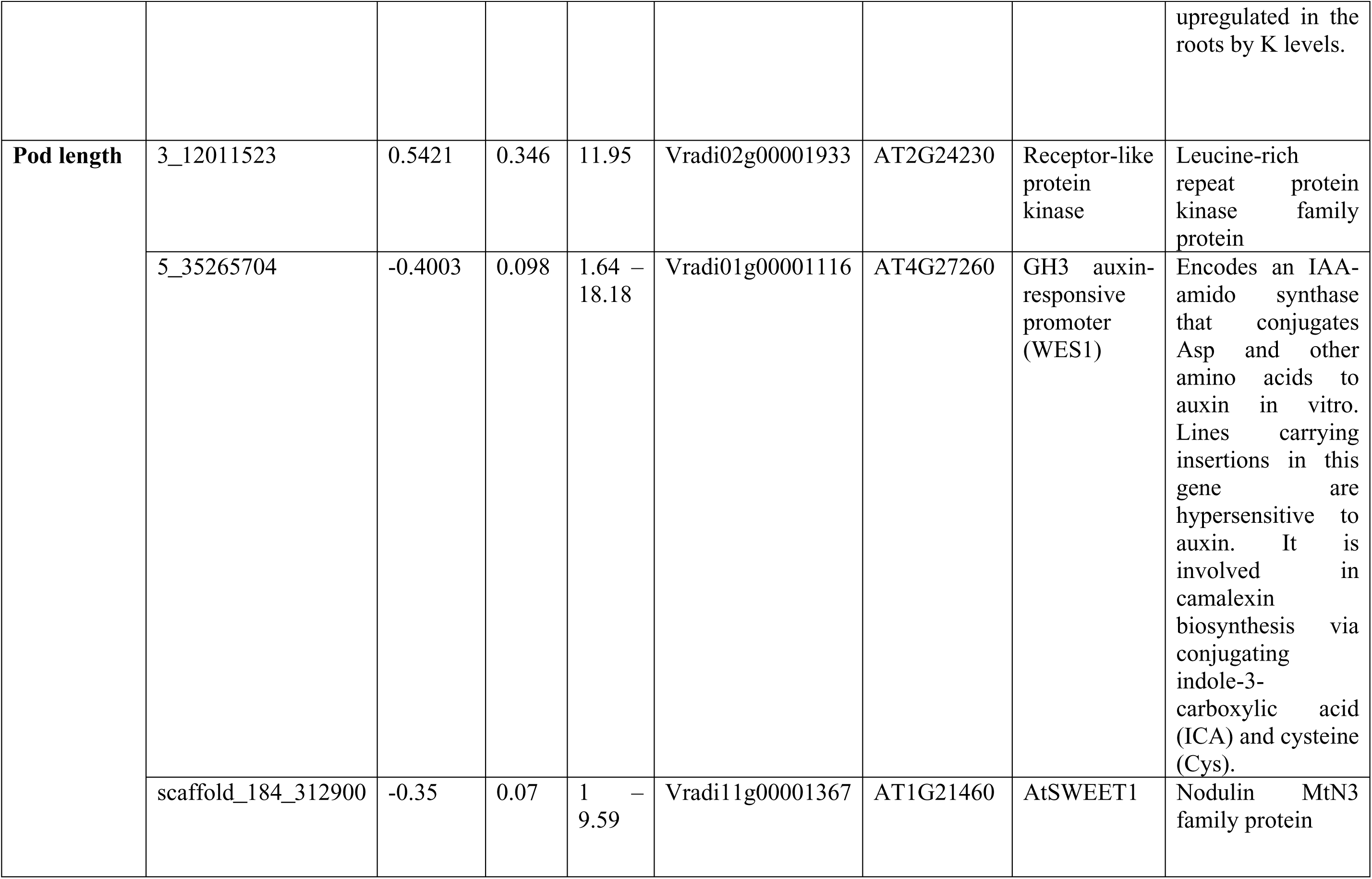
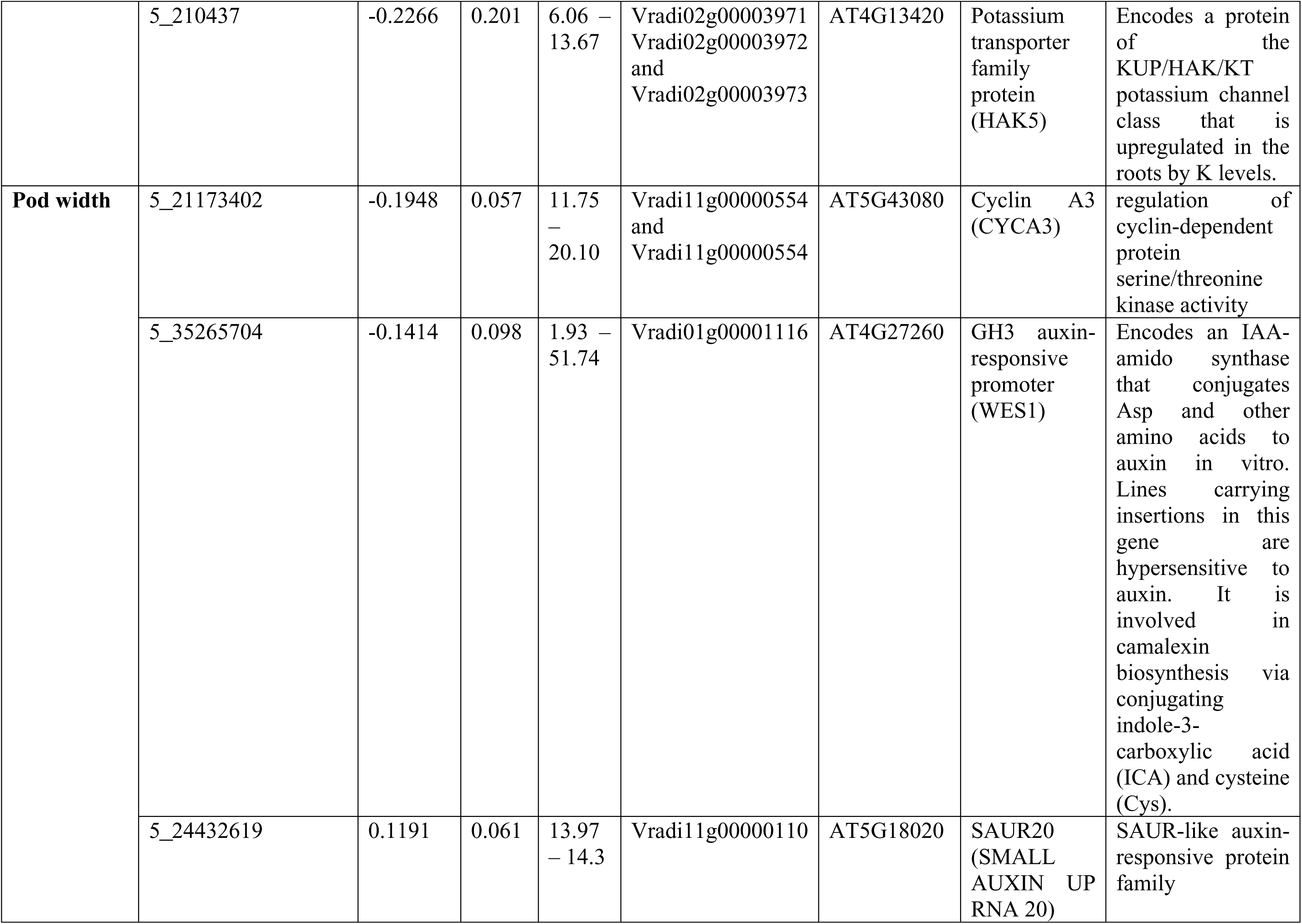
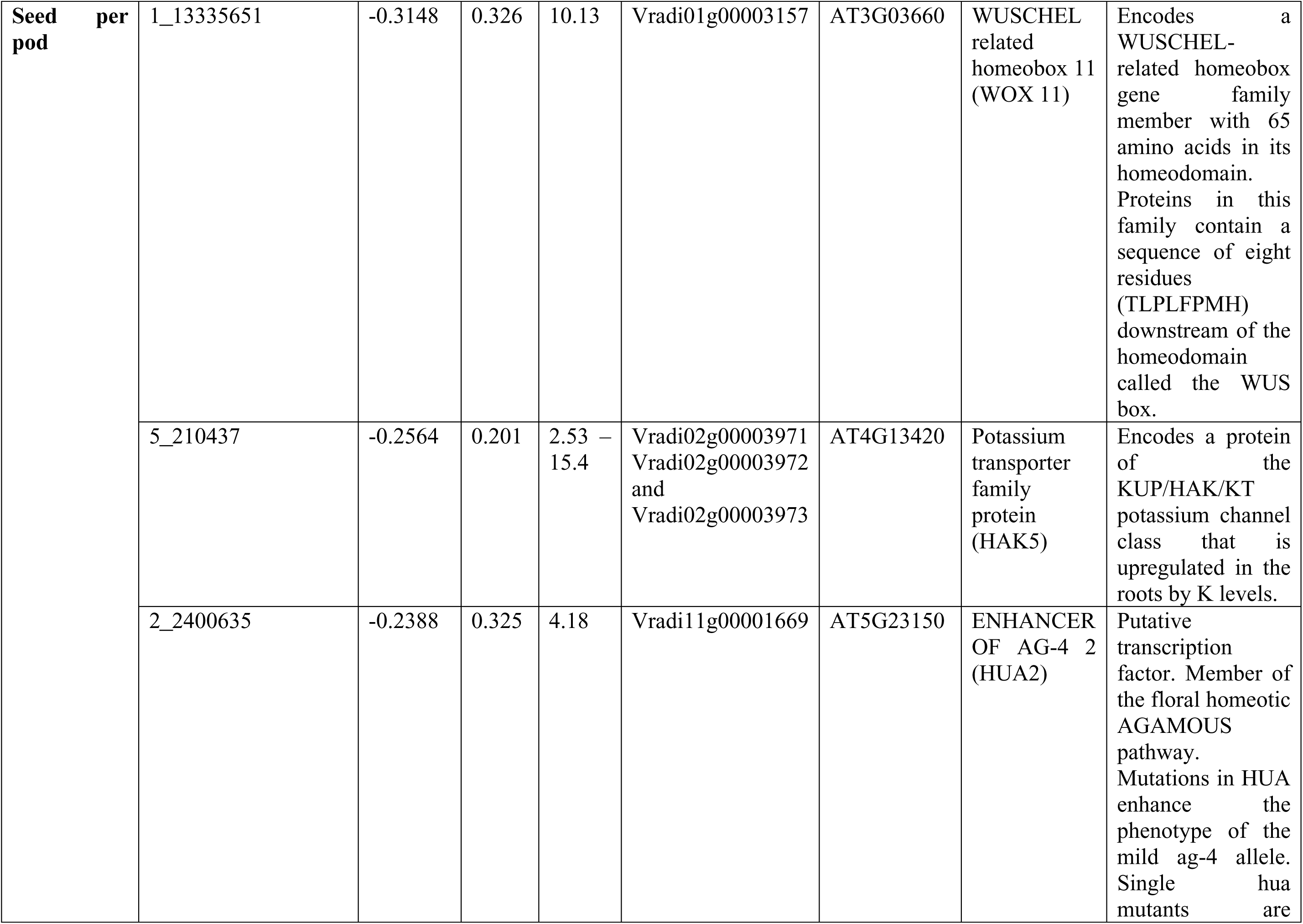
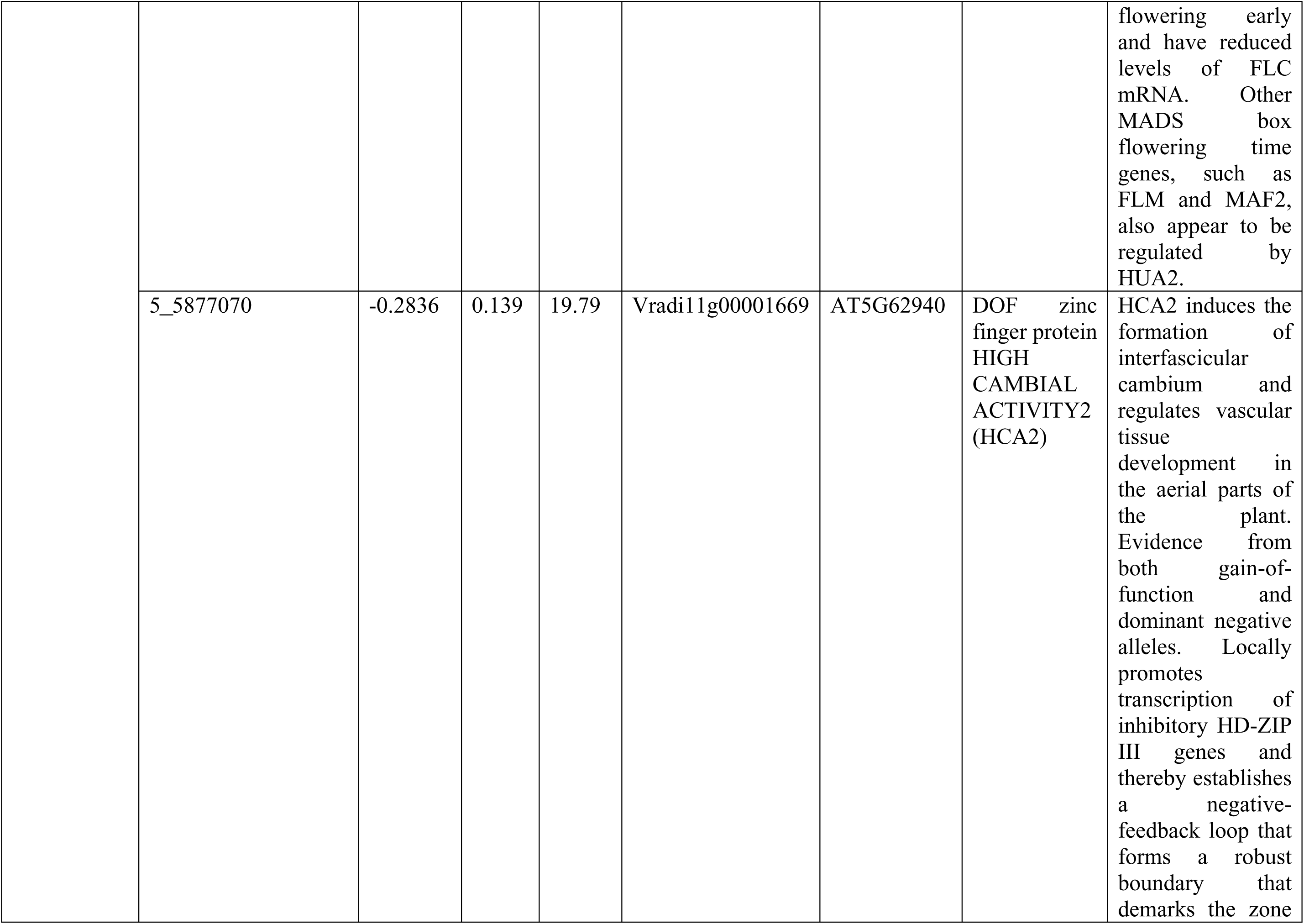
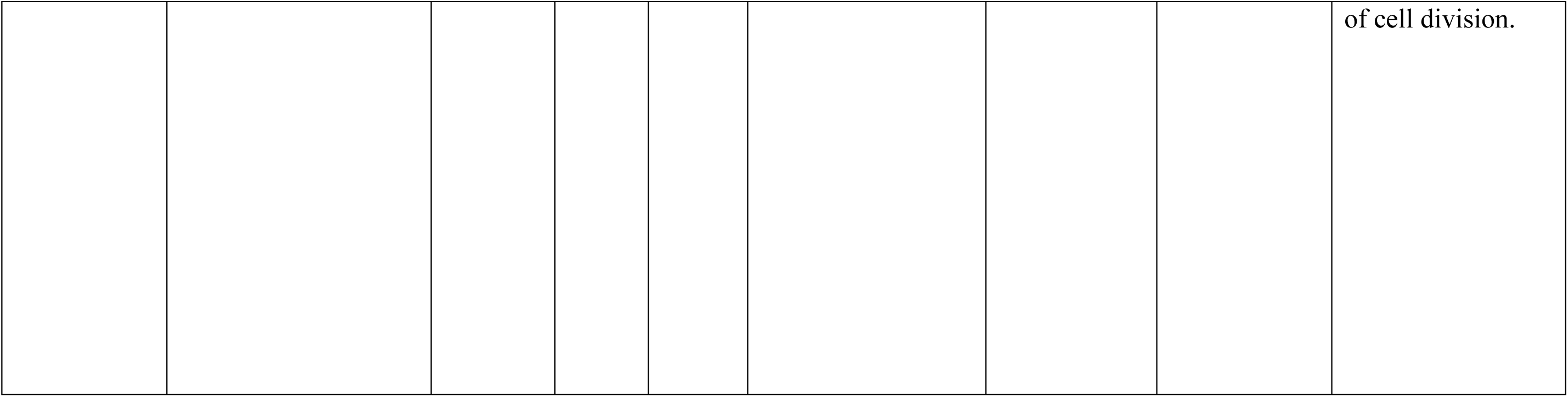
Details of identified significant marker-trait associations from the three models and the candidate genes associated with them extracted from the mungbean reference genome *Vradiata_version 7* (Sandhu and Singh 2021) at the *Legume Information System* website (https://www.legumeinfo.org/Vigna/radiata/), and the *Arabidopsis* ortholog genes and protein description from BLASTp in TAIR -Arabidopsis website(Schäffer et al. 2001) (https://www.arabidopsis.org/tools/blast/).

In an interesting finding, SNP 5_35265704 (C→T), located on chromosome 1, was commonly associated with all three pod dimensional traits: pod length, pod width, and pod curvature. It has an allelic effect of −0.34, a minor allelic frequency of 0.10, and with 18 – 49% PVE. Additionally, another SNP, 5_210437 (T→G), on chromosome 2, was found to be associated with both pod length and seeds per pod traits, with an allelic effect of −0.23 and a minor allelic frequency of 0.20 by explaining 12.30 % of phenotypic variance. These two SNPs were further analyzed for comparative mapping with related species, including cowpea (*Vigna unguiculata*), common bean (*Phaseolus lunatus*), adzuki bean (*Vigna angularis*), and soybean (*Glycine max*), as well as the model legume, *Arabidopsis thaliana* (Figure 7). A total of 31 genes were identified within the LD block (532 kb) near SNP 5_35265704 on chromosome 1, out of which *Vradi01g00001101*, *Vradi01g00001100*, *Vradi01g00001119*, and *Vradi01g00001116* could be plausible potential genes based on their gene ontology. Among these genes, *Vradi01g00001116* was recognized as a candidate responsible for pod dimensional traits due to its putative indole-3-acetic acid-amino synthetase activity. The chord diagram in Figure 7A, resulting from a comparative sequence alignment, shows that this gene has multiple high-confidence alignments with multiple genes in soybean (*Glyma06G301000, Glyma12G103500, Glyma12G197800*, *and Glyma13G304000*; 71.9-84.3% identity), followed by cowpea (*Vignaun05g223100*; 92.6% identity), adzuki bean (*Vang09g02910*; 95.9% identity) and common bean (*Phvul005G111000*; 89.9% identity), all with an E value of 0, particularly more with soybean. Furthermore, *Vradi01g00001116* shares strong homology with several *Arabidopsis* GH3 genes (*AT2G47750* and *AT4G27260*), which have an auxin-responsive promoter function that influences organ elongation, as well as pod, seed development, and hypocotyl growth. The digital expression study of this gene suggests that the gene is expressed in different plant parts, including hypocotyl, leaves, flowers, pod, and seed in *Arabidopsis thaliana* (Figure 7C).

**Figure 7:**
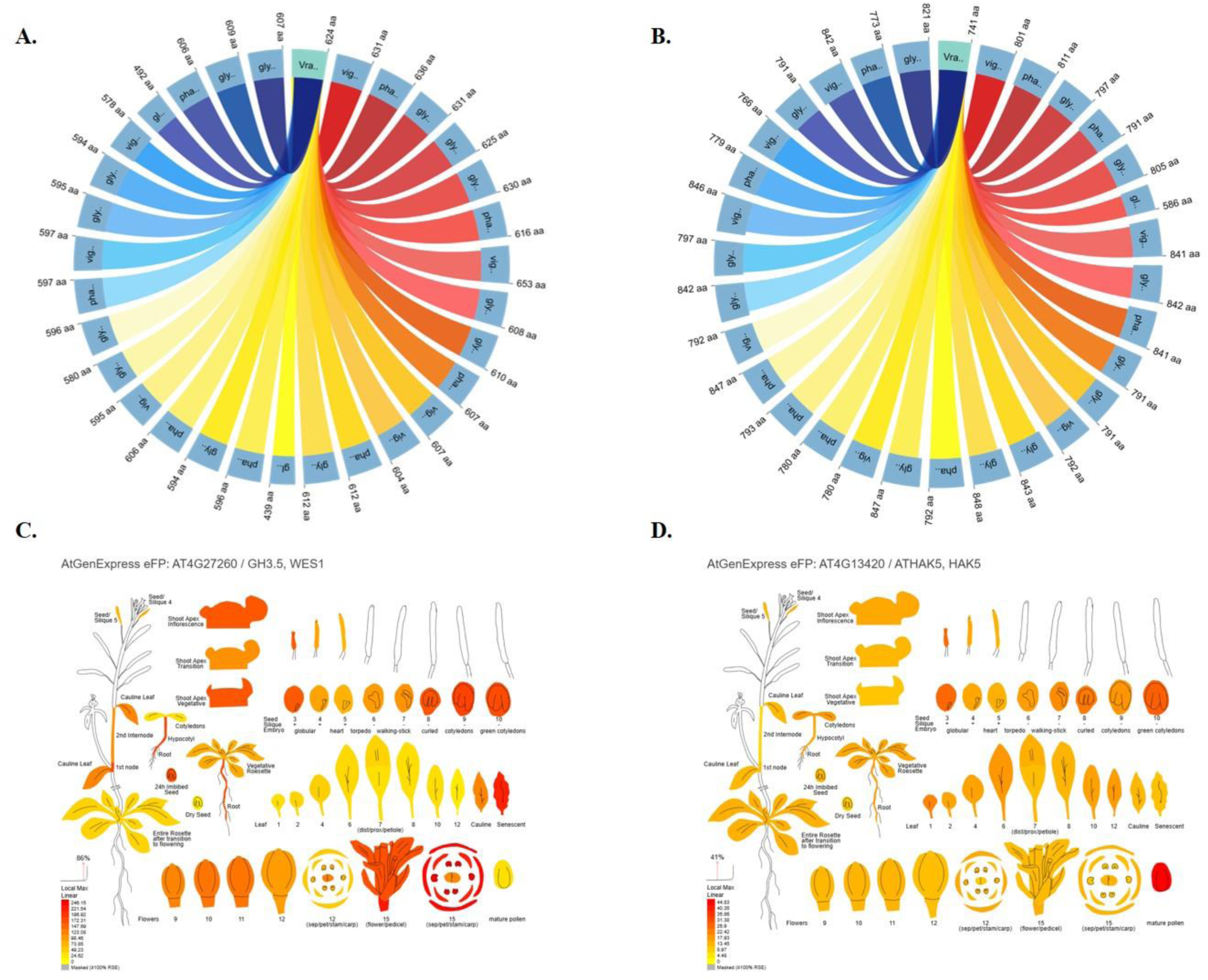
Comparative homology and tissue-specific expression of GH3 and HAK5 family genes (A, B). Chord diagrams illustrate the BLASTp-based homology of mungbean GH3 candidate proteins with orthologs from *Glycine max* (gly), *Phaseolus vulgaris* (pha), adzuki bean (Vigna angularis), and *Vigna unguiculata* (vig). Panel A corresponds to the GH3.5-like gene (*Vradi01g00001116*), while panel B corresponds to another HAK5 candidate (*Vradi02g00003971*). The thickness of the ribbons indicates the strength of the alignment and the level of sequence similarity, highlighting gene conservation across species. (C). Tissue-specific expression profiles of *Arabidopsis* GH3.5 (*AT4G27260*, WES1) and (D) Tissue-specific expression profiles of *Arabidopsis* HAK5 (*AT4G13420*) retrieved from AtGenExpress eFP browser (Klepikova et al. 2016). Expression is visualized across various developmental stages and organs, with red indicating high expression and yellow indicating moderate expression.

Another key SNP, 5_210437, located in an intergenic region on chromosome 2, was consistently associated with pod length and the seeds per pod traits across all three GWAS models and encompassing approximately 53 genes within the LD region (532 kb). Among these genes, *Vradi02g00003971*, *Vradi02g00003972*, and *Vradi02g00003973* have been shortlisted as potential candidates belonging to the potassium transporter family genes with potassium ion transmembrane transporter activity. Based on the score (bits) from the BLASTp search, Vradi02g00003971 emerged as the most promising gene among this closely clustered group. Therefore, a syntenic analysis was conducted using this gene, which resulted in multiple homologous connections with cowpea (*Vigun09g000800*; 84.5% identity), common bean (*Phvul009G262700, Phvul009G212700, Phvul009G212800, and Phvul009G212900*; 80.1% identity), and soybean (*Glyma19G263100, Glyma03G264100, and Glyma07G042500*; 77.1% identity), all with an E-value of 0 (Figure 7B). Additionally, it shows strong homology with the *HAK5* (*ATHK5*) gene family proteins in *Arabidopsis* (*AT4G13420*), which are known for their high-affinity potassium transport function. The digital expression study *in silico* revealed that this gene is expressed in mature pollen, immature pod, and seeds during the plant developmental stages in *Arabidopsis* (Figure 7D). These syntenic analysis results further support the presence of highly conserved orthologs in these species.

## 4. GENOMIC PREDICTION

Genomic prediction was conducted utilizing rrBLUPs in the rrBLUP package for four pod morphological traits using 10-fold cross-validation. The prediction accuracies, measured as Pearson correlation coefficients between predicted and observed values, varied among the traits and measurement methods, as shown in Figure 8 and Table S2. Pod width exhibited the highest genomic prediction accuracy (r = 0.852 ± 0.041), and this trait was evaluated exclusively through image-based phenotyping. Among the traits that were measured using both approaches, image-based pod length had superior predictability (r = 0.830 ± 0.042) over manual measurements (r = 0.794 ± 0.056). Similarly, image-based pod curvature showed moderate prediction accuracy (r = 0.668 ± 0.049), demonstrating a clear advantage over manual rating (r = 0.547 ± 0.061). Also, both image-based and manual-based seeds per pod had moderate prediction accuracy with no minute difference between the two methods. (image-based: r = 0.610 ± 0.089; manual: r = 0.621 ± 0.092). The observed prediction accuracies were consistent with estimates of trait heritability, supporting the genetic architecture that underlies phenotypic variation. The selection coincidence indices ranged from 0.0 to 0.4 across traits, measuring the agreement between the top 10 genotypes selected by BLUPs (Table S3) and GEBVs (Table S4). Image-based pod width, pod length (both image and manual-based), and seed per pod (manual) resulted in higher selection coincidence (0.4) with four common genotypes out of the top 10, followed by image-based seed per pod (0.3). Pod curvature trait showed poor agreement for both image-based (0.0) and manual (0.2) methods.

**Figure 8.**
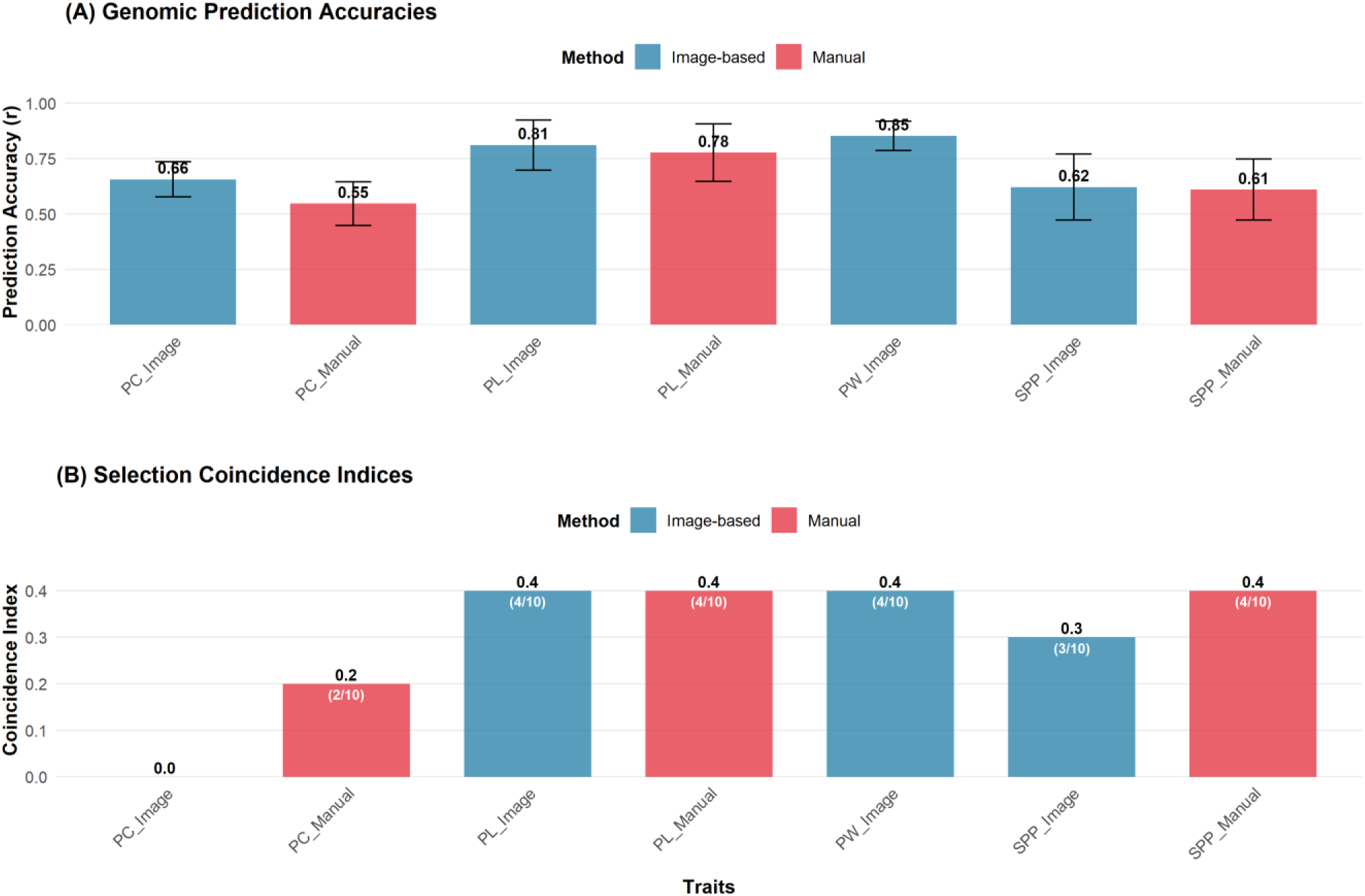
Genomic prediction accuracies and selection coincidence indices of mungbean pod morphological traits. (A). Genomic prediction accuracies (correlation coefficients, r) between observed trait BLUPs and GEBVs from 10-fold cross-validation using rrBLUP. Error bars represent standard deviation across cross-validation folds. (B). Selection coincidence indices between top 10 genotypes selected based on BLUPs versus GEBVs for each trait. Numbers on bars indicate coincidence index values, while numbers in parentheses show the count of common genotypes out of top 10 selections. (PL = Pod Length, PW = Pod Width, PC = Pod Curvature, SPP = Seeds per Pod.)

## 5. DISCUSSION

This study demonstrates the effectiveness of image-based high-throughput phenotyping in capturing subtle and complex variations in pod morphological traits of mungbean (*Vigna radiata* (L.) R. Wilczek), a crop of growing importance for sustainable protein production. By integrating deep learning-based image segmentation with genome-wide association studies and genomic prediction models, we gained valuable insights into the genetic architecture of pod morphological traits. Such knowledge can play a key role in accelerating genetic improvement and cultivar development in mungbean. The use of computer vision and machine learning has already transformed plant phenotyping by enabling high-throughput and objective measurement of complex traits (Singh et al. 2016; Singh et al. 2018; Parmley et al. 2019; Singh et al. 2021). To our knowledge, however, no previous reports have focused on image-based phenotyping of mungbean pod morphological traits.

In this study, the Epson Expression 11000XL scanner was used to get high-throughput imaging of mungbean pods with a resolution of 300 dpi. Previous studies have proven that this scanner was effective in seed and root phenotyping, showcasing its reliability in capturing detailed morphological features (Pace et al. 2014; Zhu et al. 2020; Miranda et al. 2023; Klasen et al. 2025). Our work, however, expands its application to include pod morphological traits, which involves a destructive phenotyping method. The strong correlation between image-based and manual phenotyping is particularly evident for pod length (r = 0.96) and seeds per pod (r = 0.96), highlighting the reliability of image-analysis-based trait extraction. In addition to aligning with manual measurements, the imaging pipeline allows for efficient quantification of traits such as pod curvature and width, which are typically challenging to measure at scale. By employing DeepLabV3 with a ResNet34 backbone, we achieved robust object segmentation and precise trait measurement, reducing rater bias and increasing throughput. This capability is especially important for breeding programs that manage large diversity panels, as well as for phenotyping advanced breeding lines to select potential high-yielding cultivars based on data-driven criteria, where consistency and accuracy in trait measurement are crucial.

The results reveal extensive phenotypic diversity and strong genetic control for most pod traits in mungbean, supporting their potential as selection targets in breeding programs. The observed correlations among pod dimensional parameters further indicate an opportunity to improve yield components in a coordinated manner, while the comparatively lower correlation between seed per pod and pod dimensions points to opportunities for independent selection to tailor trait combinations. The high heritability (0.74 – 0.91) of these pod morphological traits in mungbean makes them strong candidates for genome-assisted breeding. The GWAS conducted using four models (FarmCPU, BLINK, MLM, and SVEN), incorporating both manual and image-based data, identified a 60 unique SNPs out of a total of 105 SNPs associated with pod morphological traits. Several loci were found to overlap across traits, indicating potential pleiotropy or linkage that requires further confirmation. Comparative mapping, supported by GWAS results, allowed us to identify potential candidate genes at each locus on chromosomes 1 and 2. An SNP 5_35265704 is particularly noteworthy as it is consistently associated with pod length, width, and curvature. Within its LD block region (532kb), we identified a plausible candidate gene *Vradi01g00001116*, which is a member of the *GH3* family and is orthologous to *AT4G27260* in *Arabidopsis thaliana* (Luo et al. 2023). This gene has been previously studied and proven to be linked to auxin homeostasis and organ elongation. Comparative genomic analysis further revealed that *Vradi01g00001116* shares strong sequence homology (92.6%) with *Vigun05g223100* in cowpea, which resides within the major region on linkage group 1 associated with pod length (Xu et al. 2017). This syntenic correspondence suggests that the mungbean chromosome 1 locus represents an evolutionarily conserved genomic region controlling pod elongation across *Vigna* species, reinforcing the role of *GH3*-mediated auxin regulation in shaping pod morphology. Our findings reinforce earlier reports that highlight the importance of *GH3*-mediated auxin conjugation in regulating pod development and curvature in both legumes (Song et al. 2016; Lo et al. 2018; Lv et al. 2023) and *Arabidopsis*.

Another important gene, *Vradi02g00003971*, located within the linkage disequilibrium block of SNP 5_210437, is associated with both pod length and the number of seeds per pod. This gene encodes a potassium transporter that is homologous to the HAK5 family (*AT4G13420*) (Maierhofer et al. 2024). Potassium transporters regulate cell elongation and reproductive development through turgor pressure and ion homeostasis control (Nieves-Cordones and Rubio 2025). This finding suggests a direct functional link between pod elongation and the number of seeds per pod in mungbean plants. The presence of biologically relevant candidate genes highlights the potential of using GWAS outputs to identify regulatory pathways that can be directly targeted in breeding programs. Additionally, the identification of overlapping loci for pod length, width, curvature, and seed per pod suggests shared genetic regulation, allowing for the simultaneous marker-assisted selection of multiple traits.

Our results from genomic predictions demonstrate that image-based phenotyping is a viable and often superior alternative to traditional manual assessment for genomic prediction in legume pod morphology. The consistently high prediction accuracies (r > 0.60 for most trait-method combinations) indicate significant potential for implementing genomic selection in mungbean breeding programs. Pod width and pod length demonstrated the highest prediction accuracies, which corresponded to traits with significant genetic control, allowing for effective genomic selection strategies. The seeds per pod exhibited moderate heritability, resulting in intermediate prediction performance. In contrast, pod curvature showed a method-dependent ability to capture genetic signals, with image-based approaches proving more effective at quantifying heritable variation. These findings align with recent reports in other legumes and cereal crops, where high-quality automated phenotyping has improved genomic prediction accuracy compared to traditional manual measurements (Li et al. 2018; Azodi et al. 2019; Crossa et al. 2025). This supports the growing consensus that phenotyping quality, rather than just genotyping density, is a critical factor limiting the speed of breeding advancements (Araus et al. 2018).

The findings from this study can assist the global mungbean crop improvement community, especially in emerging cultivated regions like the Midwest, where there is a growing demand for plant-based protein. Genomic tools combined with high-throughput image-based phenotyping help breeders efficiently test large populations is less resource-intensive with high accuracy. Identifying true trait-associated loci and predictive markers is essential for developing effective and accurate marker-assisted selection and genomic selection strategies, which positively contribute to yield-related traits (Xiao et al. 2022; Pinto et al. 2023). In addition, genetic analysis of pod morphological traits often associated with harvest index and consumer preferences can help trait pyramid approaches. This ultimately strengthens breeders to simultaneously improve both productivity and market appeal (García-Fernández et al. 2023).

While our scanner-based imaging method has proven effective, it restricts our ability to conduct large-scale studies on trait development across different growth stages. Future research should focus on non-destructive phenotyping platforms, such as drone or rover-mounted imaging systems, which can capture pod traits over time without the need for manual harvesting. These technologies would allow us to create larger, more precise datasets while reducing the resources required for genetic analysis studies. Additionally, although our genomic prediction models have achieved moderate to high accuracy, there is still room for improvement to ensure more reliable selection decisions. The predictive power and robustness of genomic selection pipelines could be enhanced by integrating data from multiple environments, expanding the diversity of the training population, and incorporating multi-omics data.

This research demonstrates the transformative potential of combining image-based phenotyping with genomic tools in mungbean breeding. Our integrated approach enhances phenotyping accuracy, improves prediction power, and accelerates the discovery of functional genetic variants. As breeding programs increasingly adopt automated and precision techniques, such integrative approaches will be crucial for developing high-yielding, climate-resilient legume cultivars.

## 6. DATA AVAILABILITY STATEMENT

Raw phenotypic data, including both image-based and manual measurements, BLUEs, and BLUPs of these traits, are provided as supplementary files. The mungbean genotypic dataset utilized in this study is publicly accessible through the Legume Information System under accession VC1973A.gnm7.SB53 (https://data.legumeinfo.org/Vigna/radiata/genomes/VC1973A.gnm7.SB53/). All analysis scripts used in this study are publicly available at GitHub: https://github.com/vboddepalli89/Image-based-pod-phenotyping.

## 7. ACKNOWLEDGMENTS

We thank Dr. Steven B. Cannon for his constructive review of the comparative genomics and synteny analyses, which strengthened the cross-species interpretation of our results and Dr Leonardo de Azevedo Peixoto for guidance on the genomic prediction framework and validation strategy. We are also thankful to the staff in Singh labs, graduate students, and the undergraduates involved in the harvesting, imaging, data collection, and annotation.

## 8. FUNDING

This study was supported with funding from the United States Department of Agriculture-National Institute of Food and Agriculture (USDA-NIFA) Mung bean breeding #2022–67013-37120 (AS &SBC), and USDA NIFA 2023-70412-41087 (SD).

## 9. CONFLICTS OF INTEREST

The author(s) declare no conflict of interest.

## 10. AUTHOR CONTRIBUTIONS

Venkata Naresh Boddepalli: Conceptualization; formal analysis; investigation; visualization; writing-original draft; writing-review & editing. Talukder Zaki Jubery: Methodology; software; writing-review & editing. Somak Dutta: Formal analysis; methodology; writing-review & editing.Baskar Ganapathysubramanian: Conceptualization; methodology; software; writing-review & editing. Arti Singh: Conceptualization; methodology; supervision; writing-review & editing.

## Notes

https://data.legumeinfo.org/Vigna/radiata/genomes/VC1973A.gnm7.SB53/

### Competing Interest Statement

The authors have declared no competing interest.

